# RhoG, Rac1 and Cdc42 cooperation in cell protrusion revealed by multiplexed optogenetics and biosensor imaging

**DOI:** 10.64898/2026.05.12.724597

**Authors:** Frederico M. Pimenta, Jaewon Huh, Christopher M. Welch, Neha K. Pankow, Daniel J. Marston, Timothy C. Elston, Gaudenz Danuser, Klaus M. Hahn

**Affiliations:** Department of Pharmacology, University of North Carolina at Chapel Hill, Chapel Hill, NC, USA, 27599; Departments of Bioinformatics and Cell Biology, University of Texas Southwestern Medical Center, Dallas, TX, USA; VAR2 Pharmaceuticals ApS, DK-2200 Copenhagen, Denmark; The Trade Desk, New York, NY, USA 10036; Division of Otology/Neurotology, University of Michigan, Ann Arbor, MI 48109; Murty Trust, Bengaluru, Karnataka, India 560041; Duke Human Vaccine Institute, Duke University, Durham, NC, USA 27710; Department of Cell Biology, Duke University, Durham, NC, USA 27708; Institute of Human Biology, Hoffmann-LaRoche, Basel Switzerland, 75390

## Abstract

The small GTPase Rac1 controls cell protrusion for a wide variety of critical cell functions. Its regulation by upstream guanine exchange factors (GEFs) has been the focus of multiple studies, but regulation by the GTPase RhoG remains poorly understood. RhoG is known to activate the ELMO/DOCK180 GEF complex, which in turn interacts with Rac1. It is unclear which aspects of protrusion are controlled by RhoG, and which of RhoG’s effects on protrusion are mediated by Rac1. To address these questions, we developed biosensors and optogenetic tools to activate one GTPase while observing another, and to simultaneously visualize the activity of two GTPases. New tools included a photoactivable RhoG, a RhoG biosensor, and red shifted biosensors of RhoG and Rac1. RhoG and Rac1 activation events in protrusions were spatio-temporally correlated with one another and with protrusion velocity. Causal inference indicated that RhoG indeed unidirectionally activated Rac1. Photoactivation of RhoG and Rac1 indicated that specific aspects of protrusion behavior were controlled by RhoG, and only some via Rac1. Further dissection of RhoG to Rac1 signaling through simultaneous GTPase activation and biosensor visualization showed that PA-RhoG activates Rac1 predominantly through DOCK180 and that PA-RhoG can activate Cdc42 independently of Rac1.

## Introduction

Control of cell morphology is a critical aspect of cell function. Beyond its obvious importance in cell movement, development, and tissue homeostasis, morphology can modulate signaling for purposes not obviously related to cell shape, such as ruffles formed to scaffold proliferation signaling ^1^. Rho GTPases are central players in the great majority of morphodynamic behaviors. They are molecular switches that modulate morphology in response to diverse upstream regulators, integrating signals from sensors of mechanical force, membrane receptors, and proteins monitoring cell homeostasis, to name a few ^2,3,4,5^. Rho GTPases control actin polymer networks, adhesions, membrane structure, microtubules, and intermediate filaments. They interact with downstream proteins upon binding guanosine triphosphate (GTP), and are ‘activated’ by guanine exchange factors (GEFs) that accelerate release of GDP to enable GTP binding. Inactivation via GTP hydrolysis is catalyzed by GTPase activating proteins (GAPs). Guanine dissociation inhibitors (GDI) control GTPase association with the plasma membrane, where effector targets are usually found. All these regulatory events must be precisely controlled in space and time to generate cell morphology ^6^.

The GTPase Rac1, essential to cell protrusion, is among the most studied members of the Rho family. Although its activation by GEFs has been well characterized, its less canonical regulation by another GTPase, RhoG, is not well understood. Here we examine the spatio-temporal regulation of RhoG activity, and how it impacts protrusion induced by Rac1. Previous studies have shown that at the leading edge of moving cells, RhoG recruits the ELMO/DOCK180 GEF complex to focal contacts, leading to activation of Rac1 ^7,8,9^. However, it is not known which specific aspects of Rac-induced protrusion are controlled by RhoG, whether RhoG also regulates aspects of protrusion independently of Rac1, or how RhoG activation of Rac1 is controlled through subcellular localizaiton or kinetics.

RhoG is 70% homologous to Rac1. It is activated by receptors for Syndecan-4 (α_5_β_1_-integrin signaling) ^10^, EphA2 (EGFR) ^11^ and FGFR ^12^. Several upstream and downstream partners for RhoG beyond the canonical ELMO/DOCK180 GEF have been identified, including Vav2 ^13^, Vav3 ^14^, TrioGEF1 ^15^, and RhoGDI-3 ^16^, a RhoG-specific RhoGDI. RhoG has been shown to traffic along microtubules to the leading edge, potentially to mediate its effects on protrusion ^10,17,18,19^. Studies on the role of RhoG in cell motility have thus far used genetic manipulation rather than the acute effects produced by optogenetics ^9,20,21,22,23^. These studies have in some cases yielded conflicting results, perhaps because the cells could compensate for genetic manipulations. Moreover, they could not determine the time and locations of functional RhoG-Rac1 interactions in relation to protrusion events.

Here, we set out to examine the control of Rac1 by RhoG in living cells, to ask how the interaction is controlled in time and space, and which aspects of Rac-induced protrusion are governed by RhoG. It has not been possible to visualize or control RhoG activity in live cells, nor to observe Rac1 activity while manipulating RhoG. We therefore developed a fluorescent biosensor for RhoG activation and an optogenetic analog for acute stimulation of RhoG signaling. We modified the wavelengths of a known Rac1 biosensor for simultaneous imaging with RhoG, and to use together with the optogenetic RhoG. These tools enabled us to map the spatio-temporal coordination and function of RhoG-Rac1 interactions during protrusion, showing several correlated activities up to 3.9 micrometers from the cell edge. We compared the induction of protrusion by photoactivatable Rac1 (PA-Rac1) ^24^ and photoactivatable RhoG (PA-RhoG), with and without inhibition of the Rac1-GEF DOCK180, revealing that only some aspects of protrusion dynamics controlled by RhoG occur through regulation of Rac1, and supporting a role for the GTPase Cdc42 downstream of RhoG.

## Results

### A RhoG fluorescent biosensor

We developed a FRET-based RhoG biosensor based on designs used in our previous Rho family biosensors (Rac1, Cdc42, and RhoA) ^25,26,27,28,29^. The RhoG biosensor used intermolecular FRET between RhoG and the C-terminal domain of ELMO (residues 1-95), each fused to a fluorescent protein. The RhoG effector ELMO binds specifically to GTP-loaded RhoG ^8^. The C terminus of RhoG was left free for binding to guanine dissociation inhibitors (GDI), which regulate both activity and membrane localization. In RhoG’s off state there is essentially no binding between RhoG and the ELMO fragment so no FRET from the biosensor. Our primary focus was thus to optimize the extent of FRET produced by binding of the two proteins. We optimized the fluorescent proteins, the length of the ELMO peptide, and the position of the fluorophore on that peptide (Supplementary Figure S1A and S1B), finally settling on mCerulean3-RhoG and ELMO (1-95)-Ypet as the FRET pair. To verify that control of RhoG activity by upstream regulators was not affected, we used our published high content screening assay ^27,30^ (Figure 1A) to examine the potency of the guanine exchange factors (GEFs) Vav2, Dbl and Tiam1 relative to a constitutively active (CA) mutant of RhoG, and examined the ability of RacGAP1 to inactivate FRET. As controls we tested the effect of proteins that interact with other Rho GTPases but not RhoG (Rho-GDI1 and the GEF Tim). The FRET signal from the RhoG biosensor showed the expected concentration-dependent effects for these regulators. As with previous biosensors, maximizing FRET brightness enabled us to use biosensor concentrations where the biosensor acted as a tracer of RhoG behavior with minimal effects on cell behavior (Supplementary Figure S2).

**Figure 1.**
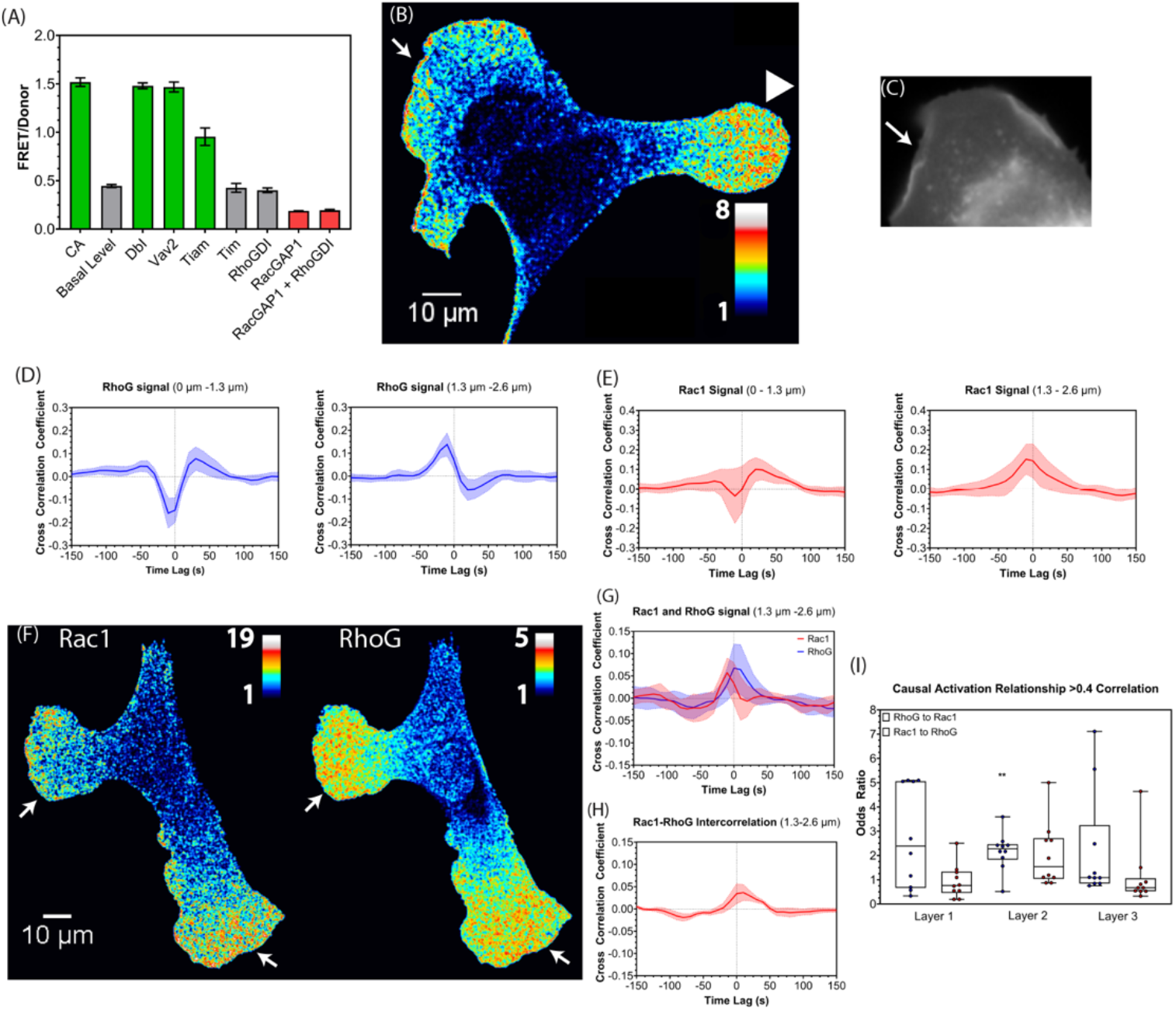
Validating the RhoG biosensor. Imaging RhoG and Rac1 activation together. (A) Effects of RhoG regulators on the FRET efficiency of the RhoG biosensor (basal =gray, activating GEFs = green, regulators not specific for RhoG = gray, activating point mutations = green; error bars SD, n=3). (B) RhoG activity in a randomly moving MEF. Pseudocolor scale indicates ratio values relative to the lowest value inside the cell perimeter (see mtl and methods). Increased RhoG activity is seen in ruffles (top left arrow) and protrusions (arrowhead). (C) Donor channel close-up of panel B highlighting ruffling regions. (D-E) Average temporal cross-correlation between RhoG biosensor activity and edge velocity, indicating the strength of association and temporal shifts between the two variables in window layers 1 (D) and 2 (E). Averages were computed over n=9 cells (see Suppl. Fig 1G for per-cell averages over all windows of a layer). (F-G) Average temporal cross-correlation between Rac1 biosensor activity and edge velocity over n =8 cells; same data representation as in (D-E). (H) Rac1 and RhoG biosensors co-expressed in MEF show activation of both GTPases in protrusions (white arrows). (I) Average cross correlation of co-expressed RhoG and Rac1 biosensor activities with edge velocity (error bands 95% CI, n=9). (J) Cross correlation between Rac1 and RhoG activity in Layer 2. (K) Odds ratio values for GTPase activation events at 0.4 correlation threshold, obtained by imaging RhoG and Rac1 biosensors in the same cell (error bars SD, n=10, Mann-Whitney non-parametric statistical test).

To control the ratio of the GTPase to the ELMO fragment in cells, these biosensor components were inserted in the same plasmid, separated by ribosomal skip sequences. Two skip sequences inserted sequentially (p2a and t2a) led to complete cleavage as assessed by SDS-PAGE ^31,32^. We created mouse embryonic fibroblast (MEF) stable cell lines expressing the plasmid under control of a doxycycline-inducible piggybac vector ^29,33^. This allowed us to minimize cell adaptation by expressing the biosensor only prior to experiments. At a variety of expression levels, cells showed the same ratio of the two biosensor chains, indicating that the skip sequences successfully controlled protein expression (Supplementary Figure 1C). We called this biosensor RhoG FLARE.dc, consistent with previous nomenclature (for fluorescence activation reporter, dual chain). We also developed a red shifted version of this biosensor using Ypet-RhoG with ELMO (1-95)-mCherry, named Red RhoG FLARE.dc (Supplementary Figure S1D-E).

Although this study focused on the functional interplay between RhoG and Rac1 during cell protrusion, we can also report that we found the RhoG biosensor associated with a perinuclear compartment, and with vesicles undergoing retrograde and anterograde transport (Supplementary Figure S3A, Movie S1). This was consistent with previous studies that showed RhoG at the Golgi apparatus and at vesicles moving along microtubules ^10,17,18,19^. In immunofluorescence experiments, RhoG colocalized with Rab4, Rab5 and Rab11 vesicles, associated with rapid endosomal recycling, but not with Rab7 vesicles, which are linked to lysosomal degradation pathways (Supplementary Figure S3B, S3C). This is consistent with previous studies indicating that RhoGDI-3 inactivates RhoG and localizes it to vesicles ^18^. Biosensor associated with vesicles was predominantly inactive (Supplementary Figure S3D). We found that RhoG colocalized with RhoGDI-3 on both vesicles and the perinuclear compartment (Supplementary Figure S3E-G). These qualitative observations confirmed that the new biosensor properly captured a variety of known RhoG behaviors, giving us confidence to move forward with more detailed, quantitative approaches that would dissect the roles of RhoG in the regulation of cell protrusion.

### Simultaneous visualization of RhoG and Rac1 activity: Relative location and timing of RhoG and Rac1 activation events during protrusion, and their causal relationship

We first examined the activation dynamics of RhoG in MEF cells undergoing random migration on fibronectin. RhoG activity was elevated in protrusions and in cell ruffles (Figure 1B, 1C, Supplementary Figure S1F, and Movie S2). The correlation of RhoG activity with edge velocity was quantified as previously described ^26,27,29^. Briefly, we generated three layers of probing windows of 1.3 µm by 1.3 µm along the cell edge. The windows maintained their distance from the edge, tracking cell edge movement. This allowed the extraction of a FRET intensity time series for each window, and for each window allowed us to correlate RhoG activity with the movement of the closest edge sector by computing the cross-correlation between FRET intensity and edge velocity at variable time lags. This indicated the strength of association between the two series, as well as potential delays of one relative to the other. The correlation functions were averaged over all non-quiescent windows in a given layer, and across multiple cells (Figure 1D and Supplementary Figure S4A).

This analysis revealed consistent spatial and temporal relationships between RhoG activation and cell edge movement. The layer closest to the edge (Layer 1, 0 -1.3 microns), displayed a strong negative correlation at lag -10s, which was due to elevated RhoG signaling in rearward moving ruffles, as well as unstable FRET signals during forward movement. A second, less significant lobe of positive correlation with a lag of +25s was associated with a decrease in RhoG signaling after the onset of retraction. In layers 2 and 3 (1.4-2.7 and 2.8-4.1 µm respectively), the correlation functions were inverted (Figure 1D and Supplementary Figures S4A). The positive peak at lag -10s indicated that, at these locations, fluctuations of RhoG activation slightly preceded the protrusive and retractive movements of the cell edge. As a corollary to this observation, RhoG activation increased already late in the retraction phase, which resulted in a secondary, weakly negative correlation lobe with a peak lag time of +25s. The same correlation structure was observed with the red shifted version of the biosensor (Supplementary Figure S4B).

This positive correlation between RhoG activation and cell protrusion was not surprising, given RhoG’s known function as an activator of Rac1 ^7,8,9^. We next sought to simultaneously image RhoG and Rac1 activity, to map their relative timing and position during protrusion, and to examine causal relationships between activation events. To accomplish this, we made a red-shifted variant of our existing Rac1 biosensor ^27^. The new biosensor used FRET between mScarlet and a Halo-Tag labeled with Janelia Fluor 669 dye (Supplementary Figure S5A-B) ^34^. It showed the same Rac1 dynamics as the original version (Supplementary Figure S5C-D). The Rac1 biosensor correlated with edge protrusion in Layers 1-3, as did the RhoG biosensor, albeit with lower statistical significance (Figure 1E, Supplementary Figure S5E). Some of our previous studies of Rac1 and edge movement in MEFs ^26^ did not indicate a negative correlation in Layer 1 and instead suggested a slightly delayed peak activation of Rac1. These quantitative differences are probably associated with the more frequent ruffling events in the present data. More recent studies from our laboratory are consistent with the correlations seen here ^27,29^.

Equipped with spectrally compatible RhoG and Rac1 biosensors, we visualized the two signaling activities in the same cell (Figure 1F, Movie S3). Expression of two biosensors produced significantly more cell perturbation than either biosensor alone. With one biosensor, protrusion and retraction dynamics mirrored control cells with no biosensor (Supplementary Figure S2). With two biosensors, even when using the minimal biosensor concentration needed for statistically significant correlations with cell edge movement, cell edge dynamics were affected (Supplementary Figure S2). Nonetheless, both biosensors retained a positive cross-correlation with edge dynamics (Figure 1G, and Supplementary Figure S6A) and they positively correlated with one another (Figure 1H, and Supplementary Figure S6B). To filter the significant noise in the multiplexed FRET imaging we computed direct correlations between Rac1 and RhoG only for windows in which both biosensor activities displayed a cross-correlation with edge velocity >0.1. In this scenario, the correlation peak was seen at +10s lag for both Layers 1 and 2, and at lag 0 for Layer 3, suggesting that RhoG activation was slightly delayed relative to Rac1, although the shifts were not statistically significant. Overall, these analyses revealed that RhoG and Rac1 are modulated concurrently during protrusion and retraction cycles.

Using the concurrent time series measurements of RhoG and Rac1 in each probing window we sought to infer causal relationships between the two signals, moving beyond mere correlation. We probed for causality using average treatment effect analysis, testing the significance of different outcomes for a treatment group relative to a control group ^35^. In the context of signal transduction, signal A is said to cause signal B when the activity of B systematically changes either upwards or downwards after A and B have interacted (treatment group), whereas B does not change without interaction of A and B. We considered interaction events between A and B as periods during which the two biosensor signals locally correlated above a threshold of 0.4 (see Materials and Methods). Of note, the periods tested were much shorter than the entire movie over which the average cross-correlation between RhoG and Rac1 peaked at values <0.1 (Figure 1H). To infer whether RhoG affected Rac1 we determined the odds for Rac1 activity to increase after a local interaction with RhoG, the odds for Rac1 activity to increase without interaction with RhoG, and then took the ratio of those values (Figure 1I). The analysis was reversed to determine whether Rac1 causally affects RhoG. Based on this computation, we found that RhoG activation at 1.3-2.6 microns from the edge caused Rac1 activation with p-values <0.01. In Layers 1+3 (0-1.3 microns and 2.6-3.9 microns respectively) the p-values were above 0.05 and thus considered not strictly significant. However, the significance systematically increased to p<0.05 when the selection of interaction events was raised to a correlation value >0.7 (Supplementary Figure 7). In the reverse direction (Rac1 to RhoG), we found weaker causal relations (p ∼0.02 at 1.3-2.6 microns from the edge, and p>0.2 in Layers 1 and 3). In contrast to RhoG to Rac1 signaling, increasing the selectivity of interaction events did not alter the significance of Rac1 to RhoG causal influence.

In summary, these biosensor and imaging studies confirmed the upstream/downstream signaling relationships between RhoG and Rac1 that had been inferred biochemically. They mapped the relative position and timing of activation events involved in this interaction, and demonstrated a causal relationship from RhoG to Rac1 activity, and not from Rac1 to RhoG.

### Photoactivatable RhoG: Different roles for RhoG and Rac1 in protrusion dynamics

After establishing a causal link from RhoG to Rac1, we wondered which aspects of Rac1 activation and protrusion could be produced by RhoG activation alone. Addressing this question required tools for acute and localized activation of RhoG. We generated an optogenetic RhoG analog that could be used together with a Rac1 biosensor, for concurrent RhoG activation, readout of Rac1 signaling, and quantitation of edge morphodynamics. RhoG and Rac1 are ∼70% homologous. Thus, we generated photoactivatable RhoG by using approaches we had applied to produce PA-Rac1 ^24^. In PA-Rac1, the photoresponsive LOV2 domain was attached to the N terminus of Rac1. In the dark, the LOV domain was tightly folded and sterically blocked Rac1’s downstream interactions. Irradiation caused unfolding of the LOV2 C-terminal alpha helix, reversibly exposing the Rac1 active site. In designing a PA-RhoG, we tested three variants, each based on attaching the LOV domain to the N terminus of constitutively active RhoG Q61L, with 0,1 or 2 amino acids removed from the N-terminus of RhoG (denoted D0, D1 and D2). Constitutively active RhoG was used so that RhoG activity would be controlled by light alone, and not by upstream regulators. In assays of co-precipitation with an ELMO fragment, the D0 variant showed the highest difference between the dark and lit states, with∼ 3x more PA-RhoG pulled down with exposure to light (Figure 2A and 2B, and Supplementary Figure S8). This was similar to the difference between constitutively active (Q61L) and dominant negative (T17N) RhoG, indicating effective control by light.

**Figure 2.**
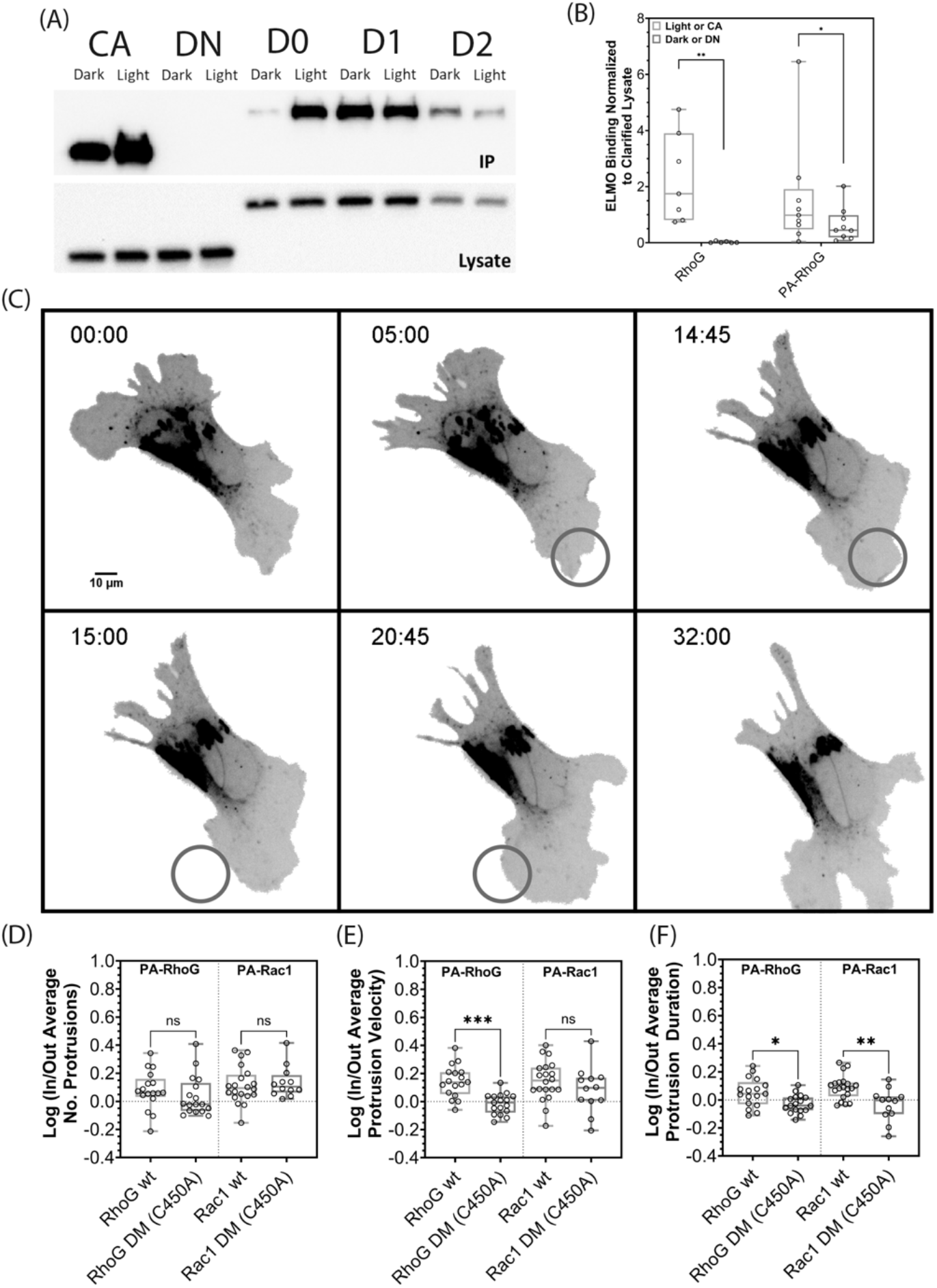
Photoactivable-RhoG (PA-RhoG) and effects of photoactivation on velocity and duration of protrusion. (A) Representative Western-Blot of immunoprecipitation under dark or light conditions of PA-RhoG constructs D0, D1 and D2 using ELMO-GST beads. Numbering represents the number of amino acids removed from the N-terminus of RhoG. CA – Constitutively Active (Q61L); DN – Dominant Negative (T17N). (B) Quantitation of PA-RhoG D1 (n=9) immunoprecipitation in dark and light conditions, and RhoG CA or DN lacking the LOV domain (n= 7) using ELMO-GST beads. (C) Representative snapshots of a MEF expressing PA-RhoG moving in the direction of the blue light irradiation. Cell was irradiated first at the lower right and then at the lower left (blue circles) (D-F) Effect of localized photoactivation of PA-Rac1 or PA-RhoG on protrusion parameters. Data normalized for cell-to-cell variation by using the log function of the ratio between the irradiated area of the cell by non-irradiated areas. LOV-wt (green, n=20 for Rac1, n=17 for RhoG), optogenetic tools with LOV-DM (red, n=12 for Rac1, n=17 for RhoG).

MEFs stably expressing PA-RhoG were exposed to a circle of irradiation of 24 µm along their edges (Figure 2C and Movie S4). The fluorescent protein iRFP720 ^36^ was fused to PA-RhoG to determine expression levels and visualize cell dynamics. PA-RhoG irradiation induced localized ruffling followed by lamellipodia formation, and in some cases cell migration, qualitatively similar to the effects of PA-Rac1 described previously ^24^. Thus, RhoG activation was sufficient to induce protrusions, leading us to ask which aspects of protrusion were produced specifically by RhoG control of Rac1.

We next asked which aspects of protrusion dynamics were controlled by Rac1, by RhoG activation of Rac1, or by RhoG acting independently of Rac1. We measured the effects of PA-Rac1, PA-RhoG or their light-insensitive C450A mutants on protrusion velocity, protrusion duration and the number of protrusions and retractions. Expression of PA-RhoG or PA-Rac1 in the absence of light did not significantly affect protrusions (Supplementary Figure S9). For experiments with acute light activation, we normalized for each cell the effects inside the irradiated circle relative to protrusion behavior in unirradiated regions, to control for cell-to-cell variability (Supplementary Figures S10, S11, see Materials and Methods).

Localized irradiation of PA-RhoG resulted in a strong increase in the velocity of protrusions (p<0.01) with a substantially less significant increase in the duration of protrusions (p<0.1) (Figure 2D-F, Supplementary Figure S11B, S11C).Localized irradiation of PA-Rac1 increased only the duration (p<0.05) and not the velocity (p>0.05) of protrusions (Figure 2D-F, Supplementary Figure S11B, S11C). Neither PA-Rac1 nor PA-RhoG had a major effect on cell retractions (Supplementary Figure S11D-F), or on the average number of protrusions (Supplementary Figure S11A). The effects of PA-RhoG and PA-Rac1 alone were consistent with previous studies indicating that RhoG can activate Rac1 ^7,8,9,37^ to produce protrusions, and that Rac1 alone is sufficient to induce protrusion. However, the differences between Rac1 and RhoG in protrusion velocity and duration suggested that RhoG activates other synergistic pathways in addition to Rac1 when it generates protrusions.

### Multiplexed photoactivation and biosensing: Synergistic RhoG Rac1 and Cdc42 signaling

Some published studies have proposed that RhoG’s action upon protrusions is mediated solely by Rac1 ^21,22^, while others have postulated that Cdc42 may also play a role ^15,20,37,38^. To address this, we observed Cdc42 or Rac1 biosensors while photoactivating RhoG. In these studies, a 24 µm diameter circle of light was used to irradiate edge segments that were morphodynamically active (Figure 3A), as previous studies ^24^ had shown that segments already active were most responsive to photoactivation. Because the cell edges were already moving, effects of GTPase activation were less striking visually, but quantitatively highly significant (Figure 3B, 3C, 3D, Movie S5).

**Figure 3.**
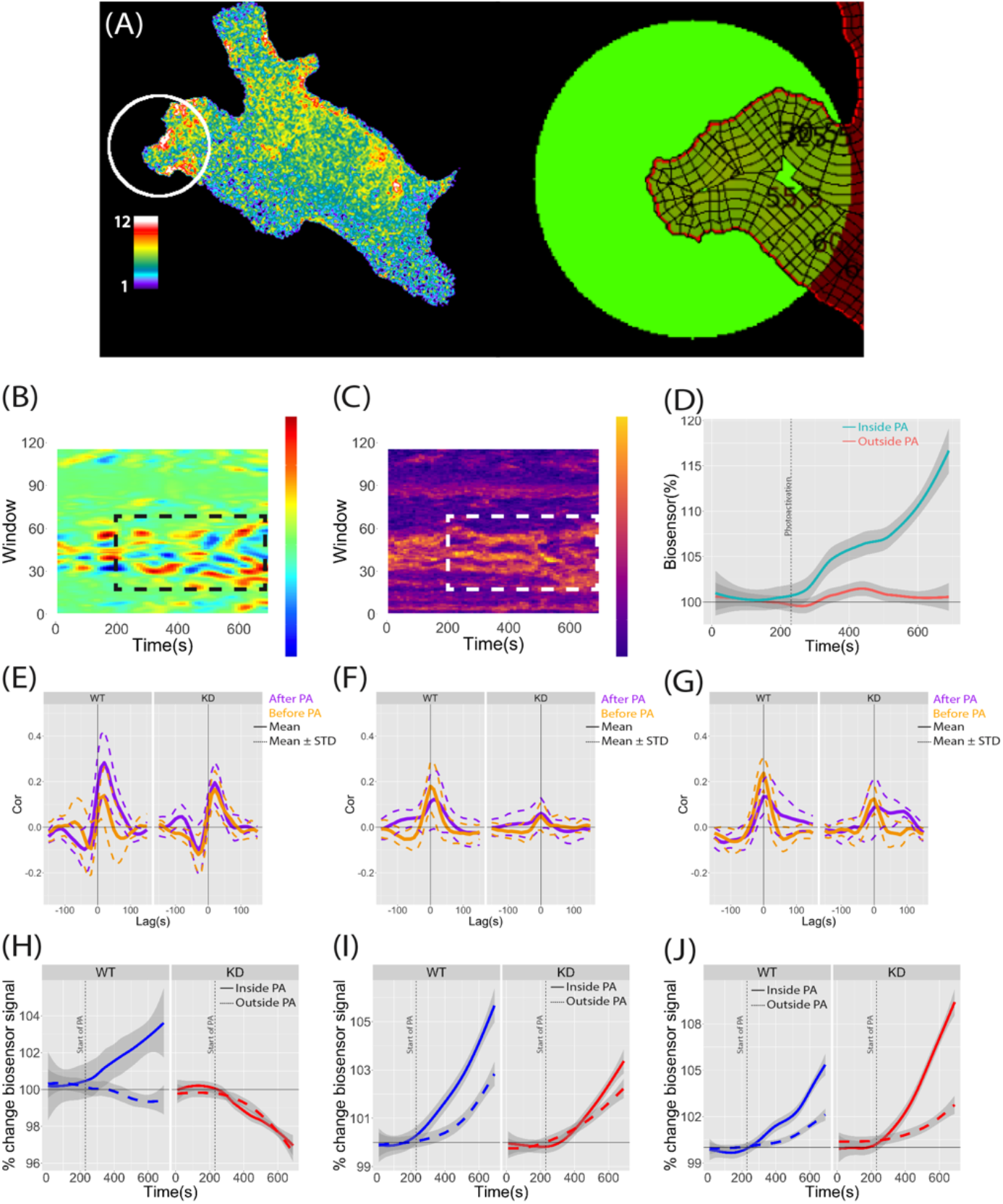
Combination of photoactivatable GTPases and GTPase biosensors reveals spatially controlled causal relationship between the GTPases. (A) Representative activity map for the Rac1 biosensor in a MEF cell expressing PA-RhoG. Red circle = area of photoactivation. Right shows windows used for analysis. Activation distal to the irradiated area was significantly decreased by reducing light intensity. Irradiation starts ∼7.5min after initiation of recording to establish a baseline for analysis (see Materials and Methods for full description). (B-C) Examples of velocity (B) and biosensor activity (C) plots as a function of elapsed time for the cell displayed in (A). Dashed rectangle shows the photoactivated region. (D) Biosensor activity as a function of elapsed time averaged for windows inside (green) and outside (red) the photoactivated region. Rac1 activation is observed after localized photoactivation (dashed horizontal line) of PA-RhoG is started. (E-G) Cross correlation plots for different combinations of optogenetic analogs and biosensors without (left) or with (right) co-expression of an shRNA for DOCK180. Using (E) as an example, photoactivation of PA-RhoG increases Rac1 correlation with edge velocity (right plots). Co-expression of an shRNA for DOCK180 abolishes this effect. (H-J) Biosensor activity as a function of elapsed time averaged for windows inside (solid lines) and outside (dashed lines) the photoactivation region. Experiments used different combinations of an optogenetic analog and a biosensor, either in the absence (blue) or presence (red) of an shRNA for DOCK180.

We computed the cross-correlation between protrusion and Rac1 activity before and after RhoG activation. Indeed, the correlation increased significantly after RhoG photoactivation (Figure 3E), indicating that activation of Rac1 by RhoG has a strong positive effect on Rac1-dependent protrusion regulation. We repeated the same experiments using Cdc42 rather than Rac1. In contrast to the above, RhoG effects on Cdc42-edge correlation were not statistically significant, and the Cdc42-edge correlation trended downwards after RhoG activation (Figure 3F). We activated PA-Rac1 in cells expressing the red-shifted Cdc42 biosensor; Cdc42-edge correlation (Figure 3G) was decreased upon Rac1 activation, indicating that Rac1 can act independently of Cdc42 activation. Together these studies indicate that RhoG can activate Rac1 to induce protrusions without involvement of Cdc42.

Biochemical evidence suggests that RhoG activates Rac1 via ELMO/DOCK180, with the tripartite complex RhoG/ELMO/Dock180 acting as a GEF for Rac1 ^7,8,9^. To test this and use Dock180 knockdown as a probe of RhoG to Rac1 signaling, we achieved robust DOX-inducible depletion of Dock180 with three different shRNA constructs (Supplementary Figure S12). We first confirmed Dock180’s role as a mediator of RhoG to Rac1 signalling by directly monitoring a Rac1 biosensor while stimulating PA-RhoG, with and without DOCK180. Knockdown of Dock180 abolished the above reported increase in correlation between Rac1 and protrusion velocity upon PA-RhoG activation (Figures 3E and 3H). Without optogenetic stimulation of RhoG, reduction of Dock180 did not affect the protrusion dynamics of MEFs (Supplementary Figure S13).

Surprisingly, depletion of Dock180 weakened the correlation between Cdc42 and protrusion dynamics overall (Figure 3F, 3G), and abolished the differences in Cdc42-edge correlation before and after photoactivation of RhoG or Rac1. This showed that Cdc42’s coordination with cell edge movement is affected by Dock180, even if not directly. This data could be explained by a Dock180-mediated activation of Cdc42 downstream of RhoG, via either RhoG-Rac1 or other intermediaries, or by the role of DOCK180 in adhesion maturation ^39,40^ and/or phosphoinositide signaling ^37,38,41,42^.

Finally, we recorded Cdc42 activity while stimulating PA-RhoG or PA-Rac1, with or without Dock180 depletion (Figure 3I, 3J). Quite unexpectedly, Cdc42 signaling increased upon photoactivation of RhoG or Rac1, regardless of Dock180 knockdown. Given the near-complete blocking of RhoG-Rac1 signal transduction by Dock180 depletion (Figure 3H), this suggested that RhoG affected Cdc42 in a Rac1-independent manner. Thus, at normal Dock180 expression levels, RhoG promotes Rac1 signaling to produce cell protrusion. At the same time, RhoG also signals to Cdc42 in a Dock180-independent manner (Figure 3I) and Rac1 activation downstream of RhoG may further contribute to Cdc42 signaling (Figure 3J). The observation that Cdc42 correlation with edge movement is weakened by depletion of Dock180 even under direct stimulation of PA-Rac1 (Fig. 3G) suggests that Cdc42 receives a third Dock180-dependent input that is Rac1-independent. All these signals to Cdc42 contribute to promotion of cell protrusion synergistic with Rac1-induced protrusions. Evidence for such synergy is further provided by our observation that PA-RhoG has a stronger effect on protrusion velocity than PA-Rac1 (Figure 2E).

## Discussion

Previous work has suggested an important role for RhoG in regulating protrusion dynamics. This work also suggested that these effects are mediated by Rac1. However, we do not know how much, and via what mechanisms, RhoG activation and effects on edge motion depend on Rac1. To address these questions we had to capture the relationship between the two GTPase and between the GTPases and edge motion directly in motile cells, requiring the generation of a new biosensor for RhoG, and red shifting Rac1 biosensors to concurrently visualize RhoG and Rac1 activity. This revealed the coordination of activation events and causal interactions between these two GTPases. Using a new optogenetic analog of RhoG together with biosensors, we were able to directly visualize the effects of local RhoG activation upon Rac1 and downstream protrusion dynamics. Consistent with previous work, this showed a direct causal link from RhoG to Rac1, but not vice versa. Of note this directionality was extracted by the implementation of a conditional average treatment effect analysis, which determined the odds ratio of effector signal increase after short term correlation of activator-effector signaling fluctuations. The correlation of RhoG and Rac1 signaling overall was weak, indicating that RhoG and Rac1 likely regulate multiple pathways. We corroborated these conclusions through direct activation of RhoG, showing that RhoG indeed activates Rac1, and that this elevated Rac1 signaling promotes edge motion, by increasing the velocity and duration of protrusions. Interestingly, direct optogenetic activation of Rac1 alone yielded mainly an increase in protrusion duration. This was likely due to the functional difference between acute activation of two signals that are in an upstream-downstream relationship. By activating the upstream signal (RhoG), regulation of the downstream signal (Rac1) by other factors was retained. In contrast, direct activation of the downstream signal overwrote this multi-factorial fine regulation. We hypothesize that RhoG’s input to Rac1 mainly controls the overall signaling strength, reflected in amplification of the protrusion strength downstream, whereas the activation and deactivation cycle of Rac1 is controlled by other inputs.

We further leveraged our ability to optogenetically activate the GTPases to show that Dock180 is a key mediator of signal transduction from RhoG to Rac1. This finding was consistent with previous biochemical analyses of RhoG-Rac1 relationships ^7,8,9^. The present causal analysis and optogenetic activation experiments indicate that the RhoG-Rac1 signal transduction is located in a narrow band of ∼1.5 um width set back from the cell edge by ∼1.5 um. In MEFs, this is the region where nascent adhesions mature to focal adhesions and where branched lamellipodial actin networks translate into a more contractile, cortical network structure. Previous reports indicated that vesicles associated with RhoG trafficking on microtubules to adhesions, where the RhoG becomes part of a tripartite RhoG/ELMO/DOCK180 complex that locally activates Rac1 ^10,17,18,19^. Our qualitative observations of the new RhoG biosensor confirmed that inactive RhoG is shuttled to peripheral adhesions on vesicles along microtubules, where it became activated. In summary, our study provided a comprehensive and quantitative dissection of a signal transduction cascade and its activity in space and time.

Equipped with an optogenetic probe for RhoG activation that was compatible with an existing Cdc42 activity sensor ^29^, we moved on to test whether RhoG was upstream of Cdc42. These experiments were motivated by controversy in earlier work, which suggested on the one hand that RhoG’s effects on morphodynamics is parallel and independent of Rac1 and Cdc42 while other studies suggested that RhoG signals are strictly upstream of Rac1 ^15,20,21,22,37,38^. The strong response of Cdc42 to RhoG photoactivation supported the notion that Cdc42 is a direct or indirect target of RhoG. However, contrary to the RhoG-Rac1 axis, the RhoG-Cdc42 axis had only minor effects on the correlation of Cdc42 activity with edge motion, indicating that RhoG’s contribution to regulating cell protrusion dynamics is mediated primarily by Rac1. Is the activation of Cdc42 by RhoG mediated by the Dock180-Rac1 axis? At first sight this seemed plausible since Dock180 depletion reduced the Cdc42 response to RhoG photo-activation. However, previous causal inference analysis harnessing a biosensor that concurrently reported Rac1, Cdc42, and RhoA activation showed that Cdc42 -> Rac1 is the predominant direction of interaction between these two signals. We therefore propose that the Cdc42 activation by PA-Rac1 is the result of a cooperative feedback activation, possibly mediated via PI3K ^38^. Somewhat surprisingly, Dock180 depletion amplified Cdc42 to Rac1 photoactivation, suggesting that this GEF has an inhibitory role in the cooperative feedback, perhaps by competition with other, PI3K-activated GEFs.

Dock180 did have a prominant role in the response of Cdc42 to RhoG photoactivation. Our interpretation is that Dock180 is essential to activate Rac1 downstream of RhoG in the first place, and that this Dock180-mediated activation of Rac1 triggers cooperativity between Rac1 and Cdc42. Consistent with this model, Cdc42 is coupled to Rac1-driven edge movements, regardless of whether RhoG is activated. However under Dock180 depletion, which abrogates RhoG -> Rac1-driven protrusion dynamics, the coordination of Cdc42 with edge movement is significantly weaker. The observation that, in the presence of normal Dock180 levels, coordination of Cdc42 with edge motion tends to decrease after photoactivation of RhoG or Rac1, aligns with our interpretation that Dock180 plays an inhibitory role on the cooperativity between Rac1, Cdc42, and cell edge dynamics. Altogether, these data indicate that RhoG signals to Rac1 via Dock180 while Cdc42 is indirectly affected by RhoG via its participation in a positive feedback between Rac1 and Cdc42. The strength of this feedback likely depends in a complex fashion on the relative expression levels of several GEFs, which may explain the differences between previous studies concerning RhoG modulation of Rac1 and Cdc42 independently or dependently. Our multiplexing of biosensor imaging and optogenetic control opened a new window into this multifactorial circuit.

Development of the new RhoG biosensor followed a now well established approach that has proven robust for several GTPase molecules ^25,26,27,28,29,43^. We identified an ELMO fragment with selective binding to RhoG’s activated conformation. Optimizing circular permutation, linker length and insertion points maximized FRET. Because the biosensor was based on intermolecular interaction between the ELMO fragment and RhoG, optimization was reduced to maximizing FRET upon GTPase-affinity reagent binding, which was accomplished through variation of linker lenghts, insertion position and circularly permuted fluorescent proteins. Importantly, the position of the inserted fluorophore preserved interaction with upstream pathways that controlled GTPase activity, and the FRET was sufficently bright that concentrations we used did not perturb normal protrusion parameters. Incorporation of red-shifted fluorophores in our published Rac1 biosensor proved relatively straightforward, and also did not change previously characterized behaviors.

Success with the new optogenetic analog of RhoG produced surprising and revealing clues potentially enabling creation of additional optogenetic GTPase analogs. The optogenetic design used here had been effective for Rac1 ^24^, but not for RhoA or Cdc42. Success with RhoG led us to study structural features of the three GTPases that might determine why only some could be succesfully ‘caged’. Previous molecular modeling studies with the optogenetic Rac1 ^24^ had suggested that Rac1 caging was effective because an FDNY sequence at aa 37-40, together with a tryptophan at position 56, led to a weak association of dark state LOV with the GTPase surface, causing LOV to sterically block the binding of downstream effectors. Irradiation caused unwinding of the LOV Ja helix, weaking this interaction and permitting the GTPase to interact with downstream molecules. RhoA lacked the aspartic acid of this motif, while Cdc42 lacked the tryptophan. Substituting tryptophan into Cdc42 produced an effective optogenetic analog. This supported our hypothesis but produced an imperfect optogenetic Cdc42 because the tryptophan had to be inserted at a position important for Cdc42 interaction with downstream proteins. RhoA lacked the aspartic acid of the FDNY motif, presenting FENY instead. Substition of D for E also produced a RhoA analog that showed strong, light dependent pulldown of RhoA effector fragments (data not shown). The substitution was again in a region important for RhoA effector binding. In contrast, RhoG was a good candidate for caging because it contained both both the FDNY and W residues seen in Rac1. We hope that sequence analysis can reveal other GTPases that will be amenable to this effective caging method.

Although we were able to use the RhoG and red-shifted Rac1 biosensors together in the same cells, this led to measurable effects on protrusion behavior. At concentrations providing sufficient signal/noise for quantitative analysis, perturbation was not observed for the individual biosesors alone. Nonetheless, we performed careful validation to ensure that the perturbations did not render our conclusions inconsistent with previous biochemical and genetic studies. In the future, especially when multiplexing biosensors for even lower abundance proteins, it will be important to design minimally perturbing molecules. We believe that much of the perturbation here comes from off target binding, where the fluorescent affinity reagent interacts with nonfluorescent, endogenous GTPase, affecting signaling without producing FRET. New methods that restrict interaction of the affinity reagent to the fluorescent GTPase analog could significantly reduce perturbation ^44^.

In summary, this paper illustrates the power of multiplexed observations and optogenetic manipulation in living cells. Only by observing or controlling two molecules together could we obtain the kinetic and spatial resolution necessary to map out the functionanally and chemically differential signaling cascades from RhoG to Rac1 and Cdc42. Challenges for using this approach in the future were uncovered, but promising leads for developing less perturbing analogs of more GTPases were also revealed.

## Materials and methods

### Molecular Cloning

Primers were purchased from Integrated DNA Technologies (IDT) and confirmed by sequencing (Genewiz). The sequences of interest were amplified by PCR from other vectors, and then assembled by restriction cloning (restriction enzymes from New England Biolabs). Commercially available kits (Qiagen and Thermo Fisher) were used for gel band purification and DNA extraction. Single point mutations for CA GTPases, and LOV lit and dark mutants, were generated by assembly PCR. Ultramers for DOCK180 were purchased from IDT, designed with restriction sites as sticky ends (XhoI at the 5’ and EcoRI at the 3’ flanks of the miR30 sequence), and annealed at a 1:1 ratio (50µM) in a water bath initially set at 98 °C and then gradually cooled to room temperature. The mixture was diluted 1:100 and ligated into the desired, restricted vector containing miRFP670 for cell labeling. iRFP720 and miRFP670 were obtained from Addgene. Other fluorescent proteins were cloned from existing vectors (see acknowledgements section).

### Cell culture and transient transfections

Mouse Embryonic Fibroblasts (MEF, ATCC XXXXXX) with a G418 TetR resistance gene, HEK-293t cells (ATCC), and Cos7 cells were cultured in Dulbecco’s Modified Eagles Medium (DMEM, Corning) supplemented with 10% FBS (Gemini) and 1 mM GlutaMAX (Corning). The cells were split every 2-3 days, at 70-80% confluency, using trypsin (Corning). Cells were only used when below passage 30. Cos7 cells were transiently transfected for bleedthrough corrections by mixing 3 µL of Fugene6 (Promega) with 100 ng of DNA for 15-30 min in OptiMEM (Gibco), which was added dropwise.

### shRNA sequences

97-mer shRNA sequences (5’→3’) were:

shRNA1

(TGCTGTTGACAGTGAGCGCCACGAAG

TTGCTTCTAAACAATAGTGAAGCCACAGATGTATTGTTTAGAAGCAACTTCGTGTTGCCTACTGCCTC GGACTTCAAGGGGTA);

shRNA2

(TGCTGTTGACAGTGAGCGACAGGTGATGTTCGAAATGATATAGTGAAGCCACAGATGTAT ATCATTTCGAACATCACCTGGTGCCTACTGCCTCGGA);

shRNA3

(TGCTGTTGACAGTGAGCGAAAAGTCGATGAT

GAAGATAAATAGTGAAGCCACAGATGTATTTATCTTCATCATCGACTTTCTGCCTACTGCCTCGGA);

and shRNA1 scrambled (TGCTGTTGACAGTGAGCGGCAACTGACCTAACCGTATATATAGTGAAGCCACAGATGTATATATACGGTTAGGTCAGTTGTTGCCTACTGCCTCGGA).

### Mouse Embryonic Fibroblasts stable cell lines

Stable, tet-off MEF cell lines were generated using the piggybac system ^33^. MEF cells were seeded in 6-well plates and allowed to reach 70-80% confluency before transfection. 5 µL of TransIT-X2 (Mirus), 500 ng of transposase (in a pUC19 vector) ^33^ and 500 ng of the piggybac vector (containing a tet recognition sequence upstream of the gene of interest and different resistance markers downstream) were incubated for 15-30 min in 100 µL of OptiMEM and then added dropwise to wells containing 2 mL of DMEM. The media was removed and replaced approximately 4 hours after addition. Cells were passed to a T-25 flask when approaching confluency; Doxycycline was added one day after transfections and the first dose of the resistance antibiotic (either puromycin, hygromycin or blasticidin depending on the selection in the desired vector) was added two days after transfection. Antibiotic concentration was doubled every 2 days - reaching a maximum at one week - and selection was maintained for a total of 2 weeks. The concentration gradients for the different selection antibiotics were as follows: (i) hygromycin (Corning): 200-600 μg/mL; (ii) blasticidin (Gold Biotechnology): 0.5-5 μg/mL; And (iii) puromycin (Gold Biotechnology) 0.5-5 μg/mL. For storage, cells were aliquoted and frozen in 90% DMEM and 10% DMSO for one week at -80°C and then moved to liquid nitrogen. Cell sorting was performed at the University of North Carolina (UNC) fluorescence activated cell sorting (FACS) Facility for all cell lines except those containing shRNAs or DOCK180-rescue, to minimize compensation in cells due to prolonged DOCK180 knockdown. In the latter, cells previously sorted for optogenetic analog expression (with or without biosensor) were always stably transfected with the piggybac-carried shRNA and selected with puromycin for 1 week before imaging.

### Biosensor Characterization

Linxe cells were seeded in 96-well plates coated with poly-lysine (3×10^5^ cells/mL) 48 hours before imaging. 24 hours prior to imaging, cells were transfected using Lipofectamine and Plus reagents (Thermo Fischer Scientific) in a 1:1:1 ratio of DNA/Lipo/plus. Cells were transfected with the different biosensor variants either alone, with constitutively active (CA), or with increasing amounts of different regulators in serum-deprived DMEM for 4 hours, after which FBS was added (10% final). CA variants of the biosensors used were: Rac1-Q61L, RhoA-Q63L and Cdc42-Q61L, RhoG-Q61L. The following Guanine nucleotide exchange factors (GEF), GTPase-activating proteins (GAP) and guanosine nucleotide dissociation inhibitors (GDI), were used: Dbl, DH-PH domain of Vav2, Tiam, Tim, Asef, p50RhoGAP, RacGAP1 and RhoGDI-1. GTPase biosensors were at 50 ng. All regulators were at 160 ng except when two regulators were combined (GDI and GAP), in which case half this amount was used for each regulator. Differences between the total amount of DNA were compensated for by adding empty vector (pTriEx) to a total of 200 ng DNA. Each assay was performed in triplicate and each individual plate contained a mock control (empty DNA carrier), plus donor and acceptor alone for bleedthrough corrections. On the imaging day cells were starved for ∼2 hours at 37 °C in Hanks’ Balanced Salt Solution (HBSS, Thermo Fischer Scientific) containing 1% FBS and 1 mM HEPES (Gibco). Plates were allowed to reach thermal equilibrium at room temperature for 30-45min. Each well was imaged at 4 different positions. Images were acquired with an ORCA-Flash4.0 V2+ sCMOS camera (Hamamatsu) coupled to an inverted IX81 Olympus epifluorescence microscope. We used Metamorph screen acquisition software (Molecular Devices), mercury arc lamp illumination and an Olympus UPLFLN U Plan Fluorite 10x objective. The following excitation and emission filters were used: (i) **RhoG mCerulean3-Ypet**: ET436/20X (Chroma) and ET500/20X (Chroma), ET470/24M (Chroma) and ET535/30M (Chroma) with a 445/505/580 ET series dichroic (Chroma); (ii) **“YFP” - “RFP” sensors**: ET500/20X and FF575/15 (Semrock) or BP-585/35 (Chroma), ET535/30M and FF01-605/15 (Semrock) or FF01-647/57 (Semrock) 445/505/580 ET series; (iii) and **Rac1 mScarlet-HaloTag**: FF01-561/14 (Semrock) and FF01-640/14 (Semrock), Dichroic ZT440/514/561/640 (Chroma), FF01-605/15 (Semrock) and FF01-692/40 (Semrock). Images were analyzed using MATLAB (The MathWorks, Natick, MA), as previously ^27,29^. Briefly, the intensity from each field of view was summed for each channel and background subtracted using values from mock-transfected wells; corrected FRET values were determined by subtracting the individual, weighted contributions of donor and acceptor channels calculated from wells transfected with only donor or acceptor respectively; FRET/Donor and FRET efficiency was then calculated for each position and averaged from at least 3 independent measurements.

### Biosensor Imaging

Biosensor expression was induced 48h prior to imaging by removing doxycycline from the media. Cells were washed, then trypsinized and seeded in a T-25 flask, and finally rewashed ∼1 hour after. On the imaging day, cells were maintained in DMEM. Approximately 2 hours prior to imaging they were seeded on coverslips coated 24 hours before with fibronectin. Cells were imaged in F-12 Ham’s medium (Caisson Labs) supplemented with 5% FBS and 1 mM HEPES. Cells were imaged using two ORCA-Flash4.0 sCMOS cameras (Hamamatsu) coupled to an inverted IX81 Olympus epifluorescence microscope, using mercury arc lamp illumination and a 40x 1.3NA Silicon oil objective (UPLSAPO40XS, Olympus). Donor channels were collected separately from FRET and acceptor channels using a two-camera adapter (TuCam, Andor) for single biosensor imaging and an image splitting W-View Gemini (Hamamatsu) coupled between the two-camera adapter and each camera for dual biosensor imaging. The following excitation and emission filter combinations were used for single and dual biosensor imaging: (i) **RhoG mCerulean3-Ypet:** FF-434/17 (Semrock) and FF01-482/35 (Semrock) for mCerulean3, FF-510/10 (Semrock) and FF01-550/49 (Semrock) for Ypet with a FF462/523 (Semrock) dichroic ;(ii) **“YFP”-”RFP” sensors**: FF01-514/3 (Semrock) and FF01-535/22 (Semrock) for Ypet, and FF01-561/4 (Semrock) and FF01-617/73 (Semrock) for mScarlet and mCherry with a ZT442/514/561m or ZT442/514/561rpc-UF2 (Chroma) dichroic; (iii) **Rac1 mScarlet-HaloTag JF669:** FF-561/14 (Semrock) and FF01-605/15 (Semrock) for mScarlet and FF-640/15 (Semrock) and FF01-692/40 (Semrock) for HaloTag JF669 with a ZT445/514/561/640 (Chroma) dichroic. Donor and FRET/Acceptor channels were separated to two cameras in single biosensor imaging using an FF509-FDi01 (Semrock) for RhoG mCerulean3-Ypet, an FF580-FDi01 (Semrock) for yellow to red sensors and an FF640-FDi01 (Semrock) for Rac1 mScarlet-HaloTag JF669. For dual biosensor imaging the same filter combinations were used for Rac1 mScarlet-HaloTag JF669 and RhoG mCerulean3-Ypet, with ZT445/514/561/640 as the dichroic, with the following modifications: Ypet emission was collected using an FF01-542/27 filter (Semrock), with channel splitting dichroics (FF640-FDi01 and FF509-FDi01) placed in the W-View Gemini and an FF580-FDi01 in the TuCam.

### Biosensor Image Analysis and bleedthrough (BT) corrections

Single cell biosensor imaging data was processed as previously described ^27,29^ with some modifications as follows: Dark current for each image was subtracted from all channels; donor channels were aligned to FRET/Acceptor channels using a registration file created from 2 µm fluorescent bead slide images collected before and after the experiments. All channels were then background subtracted and cropped to the region of interest containing the cell. Donor channels were used to generate a mask applied to all remaining channels. Bleedthrough corrections (BT) were applied to FRET. Donor, corrected FRET and acceptor were photobleach corrected. Bleedthrough contributions from donor and acceptor fluorescence to the FRET channel were determined from images of Cos7 cells transiently transfected with donor or acceptor, and imaged using the same microscope optics and illumination conditions. Bleedthrough corrections to the FRET channel were performed using the equation: *FRET*_*corrected*_ = *FRET* − *αDonor* − *βAcceptor*, where α and β are respectively donor and acceptor individual emissions in the FRET channel divided by their emission in the appropriate channel.

### PA-RhoG pulldowns and GST-ELMO beads

GST-ELMO beads were prepared following a previously established protocols ^45^. HEK-293t cells were plated in 100 mm dishes 24 hours prior to transfection. Cells were transfected with either PA-RhoG Δ1, Δ2, Δ3, RhoG Q61L or RhoG F37A at 70-80% confluency with 3.5 µg DNA, 15.75 µL Lipofectamine (Invitrogen) and 15.75 µL Plus Reagent (Invitrogen) in serum-free media, and after 4 hours supplemented with 10% FBS. For samples in dark conditions plates were wrapped in aluminum foil. 24 hours after transfection the cells were washed twice with PBS (Corning), scrapped and lysed by pipetting up and down in lysis buffer (50mM Tris-HCl (Sigma-Aldrich), 150 mM NaCl (Fisher Scientific), 1% Triton X-100 (Sigma-Aldrich) and one EDTA-free Protease Inhibitor Tablet (ThermoFisher) per 50 mL of lysis buffer, either under dark or room light conditions. The lysate was centrifuged at ∼21000 x g for 5 minutes, and the supernatant incubated with the appropriate GST-beads at 3 different concentrations of supernatant (300, 150 and 75 µL of a total of 1 mL of clarified lysate). Beads and lysate were incubated at 4°C for ∼2 h in a rocker, and afterwards washed 3 times with lysis buffer, centrifuging at ∼2300 x g for 5 min between each washing step. Beads were resuspended and boiled in a final volume of 30 µL of lysis buffer and 10 µL of a 4X LDS Buffer (Invitrogen) with 5% β-mercaptoethanol and boiled at 98°C for 30 min. Samples were run on Mini-Protean pre-cast SDS-PAGE Gels (Biorad) and imaged using a ChemiDoc™ MP Imaging System (BioRad). Proteins were then transferred to low fluorescence PVDF membranes (Biorad) using a TransBlot® Turbo™ Transfer System (BioRad) following standard manufacturer protocols. Membranes were washed 3x with Tris Buffered Saline (TBS) for 5 min and blocked for 1 hour at room temperature in a rocker with StartingBlock™ (TBS) Blocking Buffer (Thermo Scientific). Membranes were then washed 3x with TBST (0.5% Tween-20, Fisher Scientific) and incubated overnight at 4°C with StartingBlock™ Blocking Buffer T20 (TBST) Block Buffer (Thermo Scientific) and primary mouse FLAG antibody (Sigma, M2 F3165, lot# SLBL1237V). Membranes were washed 3x with TBST and then incubated at room temperature in a rocker with Blocking Buffer T20 and anti-mouse IgG secondary antibody (Sigma, A9044). Membranes were washed 6x with TBST prior to imaging to remove excess secondary antibody. Westen blot data were quantified using ImageLab from Biorad. Western blot data for clarified lysates were used to normalize for differences in total protein concentration between lit and dark mutants.

### Photoactivation

Cells were handled as described in the *Biosensor Imaging* section, except that cells were always handled either in the dark, under a red light bulb or wrapped in aluminum foil once doxycycline had been removed. Photoactivation was as described previously ^46^. Briefly, a 445 nm laser line (Cobolt, for photoactivation) and mercury arc lamp illumination (for widefield/fluorescence imaging) were combined in the back port of the microscope using a ET750sp-2p8 (Chroma) for laser reflection followed by a Longpass Dichroic Mirror, and 490 nm Cut-On (Thorlabs) to reflect lamp illumination and pass 445 nm illumination. To acquire Donor and FRET channels simultaneously in experiments using optogenetic analogs and/or knockdown (iRFP720 and miRFP670 respectively), a filter wheel was placed between the long wavelength camera and the TuCam adapter, allowing the donor channel (i.e. Ypet) to be collected in a single camera with other fluorescent proteins imaged sequentially in the long wavelength camera. The increase in distance between the long wavelength camera and the sample caused by the filter wheel was compensated for in the short-wavelength camera (i.e. donor channel) using a 1.2x lens. For experiments encompassing only optogenetic molecules, cells were imaged every 10 sec for a total 15 min, with photoactivation occurring from minutes 1 to 11; for optogenetic stimulation and biosensor imaging, cells were imaged every 10 sec for a total 15 min, with photoactivation from minutes 7.5 to 15; For both, a circle with a diameter of 24 µm was raster-scanned for 500 ms every minute during the photoactivation period, with a total power of 30 µW measured before the objective. iRFP720 was imaged using FF01-700/13 (Semrock) and FF01-735/28 (Semrock) as excitation and emission filters respectively, and FF705-Di01 (Semrock) as dichroic; miRFP670 was imaged using FF01-635/18 (Semrock) and FF01-673/11 (Semrock) as excitation and emission filters respectively, and ZT445/514/561/640 as dichroic. The miRFP670 fluorescence was only collected in the first and last frame of each movie, and iRFP720 was collected for the full duration of the movie for optogenetic experiments, and only in the first and last frame of each movie for experiments with biosensors. Optogenetically-compatible biosensors were imaged using the optics described in the *Biosensor Imaging* section.

### Quantification of the Effect of Photoactivation of PA-Rac1 and PA-RhoG on cell edge dynamics

MEFs stably expressing one of four optogenetic tools – PA-RhoG, PA-Rac1 or their light-insensitive variants (dark mutant, DM) - were locally irradiated with blue light for 500 ms every 10 sec for a total of 10 min. The irradiation area was defined by a circle with 24 µm of diameter. We first compared the effect of localized photoactivation in protrusion and retraction events by analyzing proximal and distal regions to the photoactivation area (Figure S9). Parts of the cell were considered proximal (or inside the photoactivation region) if within a 346 µm^2^ area from the center of the irradiation spot. Regions outside this area were considered distal (or outside the photoactivation region). PA-Rac1 increased both the velocity and duration of protrusions when comparing proximal and distal regions to the photoactivation area, while PA-RhoG only increased the velocity of protrusions. The duration of retractions in areas proximal to the photoactivation region decreased for PA-Rac1. During our experiments, we empirically determined that PA-Rac1 and PA-RhoG only effected changes in cell edge dynamics when choosing regions prone to protrusion. In other words, if we chose areas with a retractive behavior, photoactivation of PA-Rac1 or PA-RhoG would not induce protrusions. To account for this, we next compared the photoactivated and non-photoactivated regions using the light unresponsive LOV mutant (DM). The effects observed for PA-Rac1 and PA-RhoG when comparing proximal and distal regions were not observed when comparing the proximal regions of PA-Rac1 or PA-RhoG and their respective light-insensitive mutants. We instead saw that our selection bias resulted in regions that protruded more even for cases with the light-insensitive mutants. The lack of a statistically significant response for PA-Rac1 was puzzling given our earlier reports. We hypothesized this could be due to inter-cell variability of the expression level of the optogenetic tool or basal level of cell edge dynamics. We corrected this by normalizing each cell by dividing proximal and distal regions (Figure S10). We then normalized across cells by using a log function. Comparing each optogenetic tool with its light-insensitive counterpart showed that photoactivation of PA-Rac1 increases the duration of protrusions. Photoactivation of PA-RhoG on the other hand increased the duration and the velocity of protrusions.

### DOCK180 Knockdown

MEFs stably expressing the different constructs (shRNA, scrambled or mock) were generated as previously described ^29,33^. Doxycycline was removed and cells plated in 100 mm dishes and allowed to express the different constructs for 0, 24, 48 or 72 hours. Prior to lysis, cells were imaged to confirm miRFP670 expression and washed three times with dPBS (Gibco). Cells were then lysed in 1X LDS Buffer (Invitrogen) with 1% β-mercaptoethanol and boiled at 98°C for 30 min. Imaging, SDS page, transfer and Western Blotting were performed as described in the *PA-RhoG pulldowns and GST-ELMO beads* sections with small modifications: membranes were cut for separate overnight incubation at 4°C with either DOCK180 (C4C12) or GADPH (14C10) primary antibodies (Cell Signaling) at 1:1000 dilution, and incubation with a secondary anti-rabbit IgG (H+L) DyLight™ 800 4X PEG Conjugate antibody (Cell Signaling).

#### Time series analysis

To study dynamic subcellular activities relative to edge motion, we computationally tracked the cell boundary movement over time and subsequently defined a cell-shape invariant coordinate system allowing registration of movement and signaling. Cell boundaries were segmented using intensity thresholding of the Donor channel. Cell boundary velocities were derived from pixel-by-pixel matching of cell contours between consecutive time points ^47^, as described in Cai et al. 2014.

Upon definition of the cell edge motion, the segmented cell masks were partitioned into sampling windows of size 4×4 pixels for 330nm pixels and 5×5 pixels for 260 nm pixels matching a 1.3 µm × 1.3 µm for single windows. One of the windows within the outermost layer at the first time point was set to be the origin of the window grid. Window locations were propagated through the time frames using the boundary displacements.

The software for cell edge segmentation, tracking, and sampling window definition can be downloaded from https://github.com/DanuserLab/u-register, along with further documentation of the algorithms in https://doi.org/10.5281/zenodo.15765708.

### Cross-Correlational analysis of edge velocity and biosensor activity

#### Hidden Markov modeling to determine states of protrusion and retraction

To identify the temporal behaviors of edge velocity maps, we applied Hidden Markov modeling as implemented in the R package ‘depmixS4’ ^48^. The modeling requires a priori selection of the number of hidden states. The model output assigns each time point of the series to one of the hidden states and determines average and variance of the data within each state. For consistent output the numerical seed is initialized prior to the model fitting by the function set.seed().

To guarantee consistent state selection across different cells, we concatenated the velocity time series of all cells into a single long vector with missing values inserted between the series. The data is saved as a depmix object and computed using the fit() function without setting initial transition probability. Models were computed with an increasing number of states until the minimum proportion of a single state reached 5%. For the majority of the presented analyses this criterion yielded 8 states. Four of the states had positive average velocity and thus defined protrusion events. The remaining four states with negative average velocity defined retraction events.

### Protrusion/Retraction analysis

All statistical comparisons of protrusion and retraction dynamics between experimental conditions were performed on the basis of each cell defining an elementary experimental unit. That is, cell motion metrics were averaged over all windows of a cell and then entered as one data point into the cell-to-cell distribution over that metric.

Protrusion initiation frequency was analyzed by counting the total protrusion events in a window over the duration of entire movie. Protrusion events comprise an uninterrupted sequence of time points assigned to a protrusion state. To avoid undue influence of spurious events we included only sequences longer than 40 seconds. Events fulfilling this criterion were then also used to compute the average protrusion velocity and event duration for a cell. For all three metrics we applied the non-parametric Wilcox rank sum test to the cell-to-cell distributions in order to determine significant shifts in protrusion behavior between experimental conditions.

The same analysis was performed for retraction events, which comprise an uninterrupted sequence of at least 40s duration assigned to a retraction state.

### Causal Analysis

We inferred the level of causal influence from one signal to another, say A to B, by calculating the odds that B changes its activity after an interaction with A over B changing its activity without interaction. Relations A to B with an odds ratio significantly above 1 indicate in this framework that the activity of signal A has a causal influence on the activity of signal B. Importantly, signal B may very well be causally influenced by further unobserved inputs.

This analysis critically relies on the definition of interaction events. We considered time periods with highly correlated fluctuations between RhoG and Rac1 in the same probing window as periods where the two signals may be functionally interacting. To identify periods of local correlation between RhoG and Rac1, we used a Hidden Markov Model to determine interaction states defined by a pairing of intercept and slope in a linear regression model between the activity of signal A (effector) and the activity of signal B (target). The predetermined number of interaction states was 6. We ordered the states by decreasing average target signal activity level during the putative interaction. For the definition of interaction events, we then utilized HMM states 3 and 4 with a target signal close to 0 in Z-scores. With this choice, interaction events could yield either an increased target signal compared to the level before the event, or the target signal would stay flat or decrease. The correlation values vary depending on the duration of the interaction event. Short interaction events with high correlation can be a measurement error and long interaction events with low correlation can still show statistical significance. We tested the statistical significance of the correlation value by Student’s t-test and applied false discovery rate control for the p-values measured within each cell (using the function ‘qvalue’ within R) ^49^.

To measure the directional influence of signal A on signal B we divided the interaction events into two groups: Group 1 encompassed events with significant correlation above 0.4. Group 2 encompassed events with a correlation level in the interval [-0.4, 0.4]. The threshold of 0.4 was chosen such that groups 1 and 2 had approximately the same number of events, however, we performed sweeps of this threshold setting to examine the robustness of our conclusions to this choice. Given the two groups, we then calculated the odds of Group 1 events to yield more frequently an activation, i.e. an increase in signal B, than Group 2 events. These calculations were performed on a per-cell basis. The statistical significance was then derived by application of a natural log transformation of the odds ratios over all cells under the null hypothesis baseline as 0. This was then followed by a Student’s t-test for comparison of the log-transformed odds ratio.

### Optogenetically-induced Biosensor Activity Measurements

To determine the effect of optogenetic activation of signal A on the activation of signal B we computed for each sampling window layer the intersection between the activating light spot and the window grid to determine the window indices inside and outside the photoactivation region. Next, we normalized per cell, layer, and photoactivation state (i.e. inside or outside the activation area) the activity values to the overall average value near the photo activation timing, which is frame 22, 23, 24. This results in a single line of activity per cell/layer/photoactivation that has value near 100% at the moment of photo activation. We could then compute time courses of relative change in biosensor FRET ratio integrated over all windows inside the photoactivation region despite signaling gradients between layers. Normalization also allowed us to merge the data from multiple cells into average time courses despite their substantially differing levels of baseline signaling and differing morphologies affecting the signaling gradients. We noticed for most experiments that photoactivation had also an effect on the regions outside the light spot. This outcome is expected as activated signaling molecules diffuse and potentially also trigger signaling cascades with spatial propagation. In contrast, for the condition of RhoG activation with Rac1 monitoring under knock-down of Dock180, we observed an equally decreasing signaling curve both inside and outside the activation area. This outcome is likely caused by photobleaching. As our model suggests, under Dock180 depletion, RhoG activation has little effect on Rac1 signaling, resulting in higher penetrance of bleaching effects. Overall, interpretation of these data have to be limited to the divergence of the time courses collected inside vs outside the photoactivation spot.

## Supporting information

Movie S1. RhoG cellular distribution

Movie S2. RhoG spatio-temporal activity

Movie S3. Imaging the activities of Rac1 and RhoG simultaneously

Movie S4. Photoactivation of PA-RhoG is sufficient to induce protrusions

Movie S5. Photoactivation of PA-RhoG while imaging Rac1

## Acknowledgments

The authors thank M. Calabrese (University of North Carolina at Chapel Hill) for constructs containing piggybac vectors; J. Sondek and K. Ravichandran (University of North Carolina School of Medicine and the University of Virginia School of Medicine) for ELMO and DOCK180 expression plasmids; R. Tsien (University of California San Diego) for the mCherry expression plasmid; and M. Ehlers (Duke University) for EGFP-Rab4, EGFP-Rab5, EGFP-Rab7, and EGFP-Rab11 expression constructs. This project was supported by grants from the National Institutes of Health to K.M.H. (R35-GM122596) and T.E. (R35GM127145). J.H. was supported by the Human Frontiers in Science Program fellowship LT000911/2018-C and C.M.W. was supported by fellowships from the NIH (T32 GM008719, and F30HL094020-02).

## Author contributions

FP performed the bulk of cell biology experiments and analysis, with studies of RhoG in vesicle trafficking carried out by CW. NK developed PA-RhoG. CW and FP developed the RhoG biosensors. DM developed the color-shifted Rac1 biosensor and supported cell studies. TE helped with analysis of RhoG behavior on vesicles. JH carried out analysis of causal relationships and the bulk of analysis in Figure 4, with help from FP. KH, GD, FP and CW conceived the study, which was directed by KH and GD. FP, GD and KH wrote the paper with input from all.

