## Supplemental Information and Movie Legends for "RhoG, Rac1 and Cdc42 cooperation in cell protrusion revealed by multiplexed optogenetics and biosensor imaging"

### Supplementary Figures

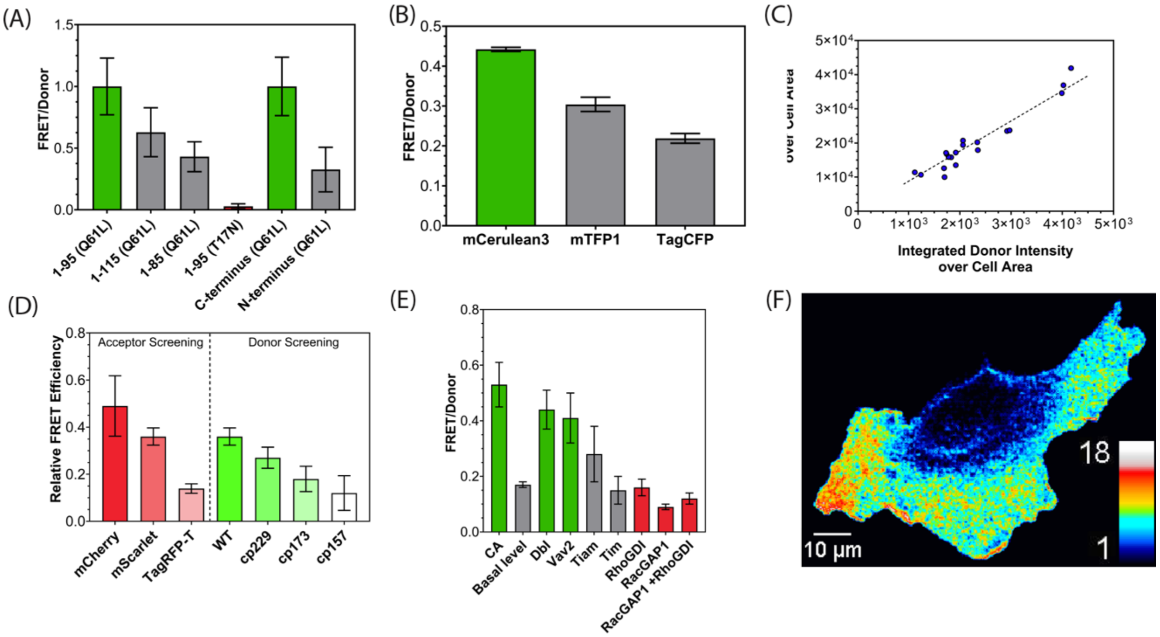


**Fig. S1. FRET biosensors for RhoG. (A)** Optimization of the dual-chain RhoG FRET biosensor by varying the length of the ELMO fragment and the position of the fluorescent protein (N- or C-terminus). RhoG with constitutively active (CA, Q61L) or dominant negative (DN, T17N) mutations were compared to maximize the FRET/Donor ratio. (Error bars SD, n=3). **(B)** Optimization for dual-chain RhoG FRET biosensor using Ypet as acceptor (SD, n=3). **(C)** Plot of Donor (mCerulean3) and Acceptor (Ypet) expression for MEFs stably expressing the dual-chain FRET RhoG biosensor. The ratio of donor to acceptor remains constant across a range of expression levels. Each dot represents an individual cell. **(D)** Acceptor and donor optimization for the red-shifted dual-chain RhoG FRET biosensor, using different acceptor and donor combinations. Donor and acceptor positions were identical to those in the version described above. Different acceptors were tested with Ypet as donor. Different donors were tested with mScarlet as the Acceptor. cp stands for circular permutants, and the number denotes the aa where the circular permutation was introduced. The Ypet-mCherry combination showed the highest FRET efficiency. **(E)** FRET efficiency of red-shifted dual chain RhoG FRET biosensor alone (Basal Level, gray) or when exposed to upstream activators (green), inhibitors (red), upstream regulators not specific for the tested GTPase (gray), and activating point mutations (green, CA) (error bars SD, n = 3). **(F)** RhoG activity imaged using the RhoG Ypet-Cherry sensor in a randomly moving MEF.

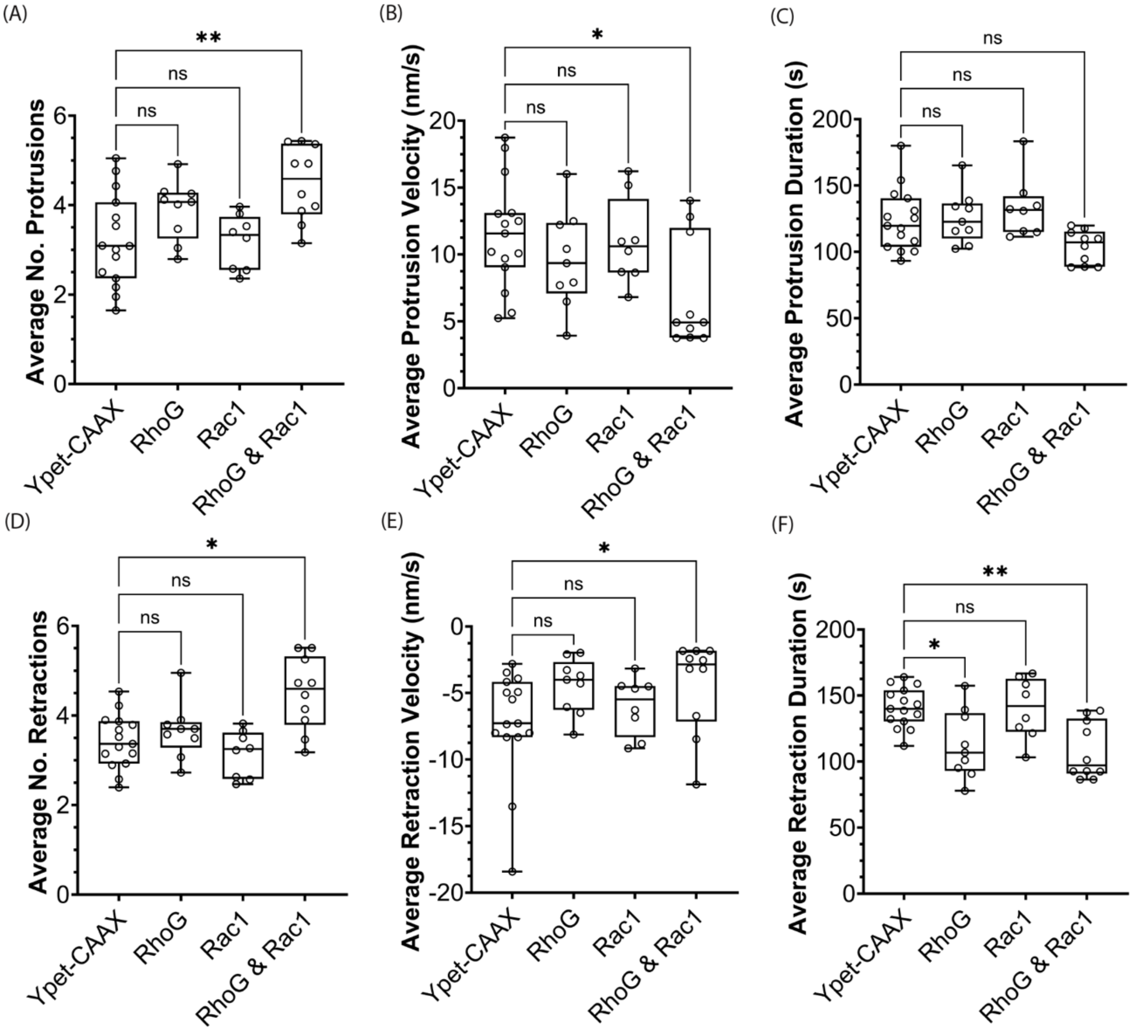


**Fig. S2. Effect of biosensor expression on cell edge dynamics.** MEFs expressing Ypet-CAAX (n=15), the RhoG biosensor based on mCerulean 3 and Ypet (n=9), the Rac1 biosensor (n=8), or both (n=10). Comparison of average frequency (A and D), average velocity (B and E) and average duration (C and F) of protrusions (A-C) and retractions (D-F) for the control group (Ypet-CAAX) and biosensors. Dunn's Test, one-way ANOVA non-parametric test (Kruskal-Wallis) used for statistical analysis, n.s. – $p > 0.05$, * - $p < 0.05$, ** - $p < 0.01$.

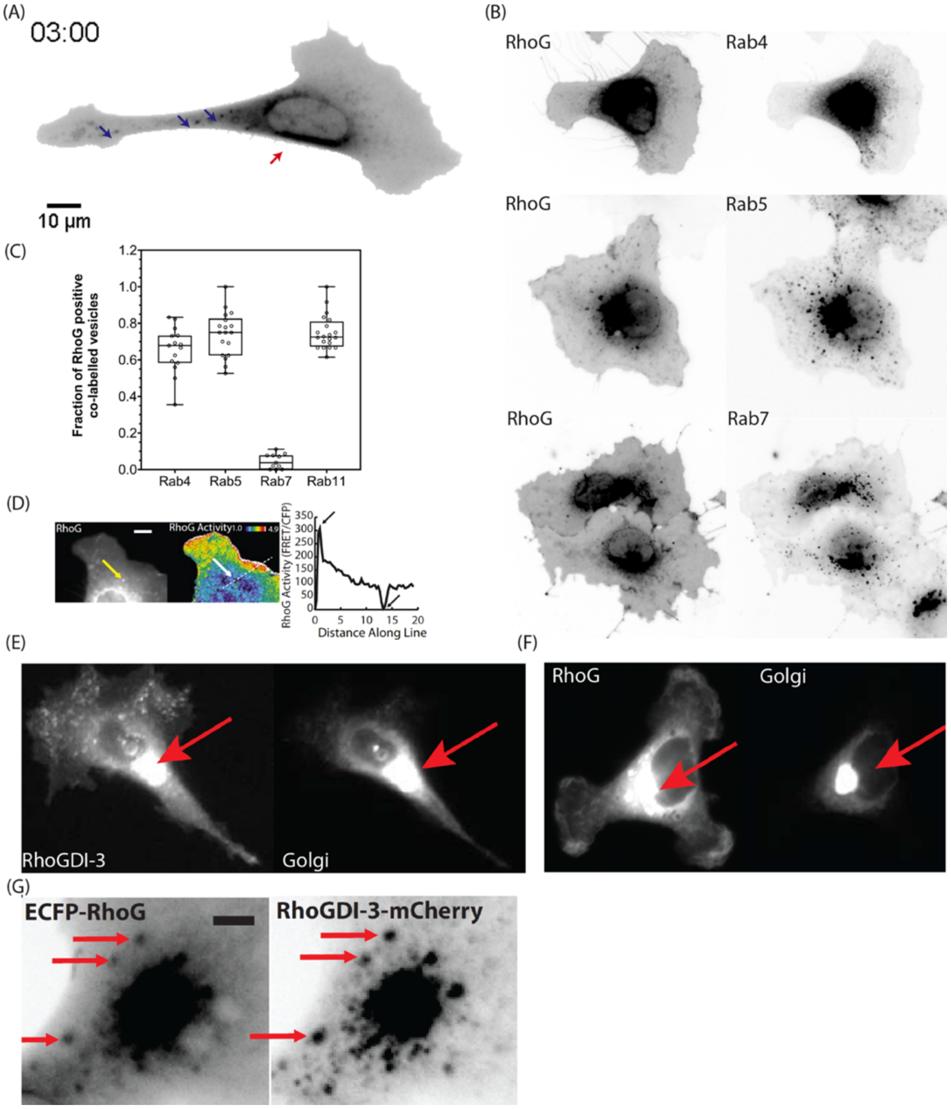


**Fig. S3. RhoG co-localization with intracellular markers and with RhoGDI-3.** (A) Representative image from a MEF expressing the RhoG biosensor. Vesicles (blue arrows) bearing RhoG move to and from the perinuclear region (red arrow). Related to Movie S2. (B) Widefield microscopy images of live Cos-7 cells transiently transfected with CyPet-RhoG and mCherry-tagged Rab4, Rab5 or Rab 7. RhoG-loaded vesicles colocalize with Rab4 and Rab5 but not with Rab 7. (C) Quantification of the fraction of RhoG-positive vesicles co-labelled with Rab4, 5, 7 and 11 (SEM, n>11 cells, n > 270 vesicles). (D) Visualization and quantitation of RhoG activity at vesicles (arrow points to same vesicle in the CyPet-RhoG image at left and the RhoG biosensor image at right) via FRET imaging. Line scan illustrates positive activity at the cell edge and activity below background at vesicles. Scale bar, 5 μm. Activity scale is indicated by heat map in upper right corner of the figure. (E-F) The perinuclear localization of RhoG and RhoGDI-3 at the Golgi apparatus in a MEF. Co-expression of (D) mVenus-RhoGDI-3 as a marker for the Golgi apparatus and (E) mCherry-RhoG as a marker for the Golgi apparatus. (G) Close up view showing co-localization of RhoG and RhoGDI-3 in vesicles (scale bar, 20 μm).

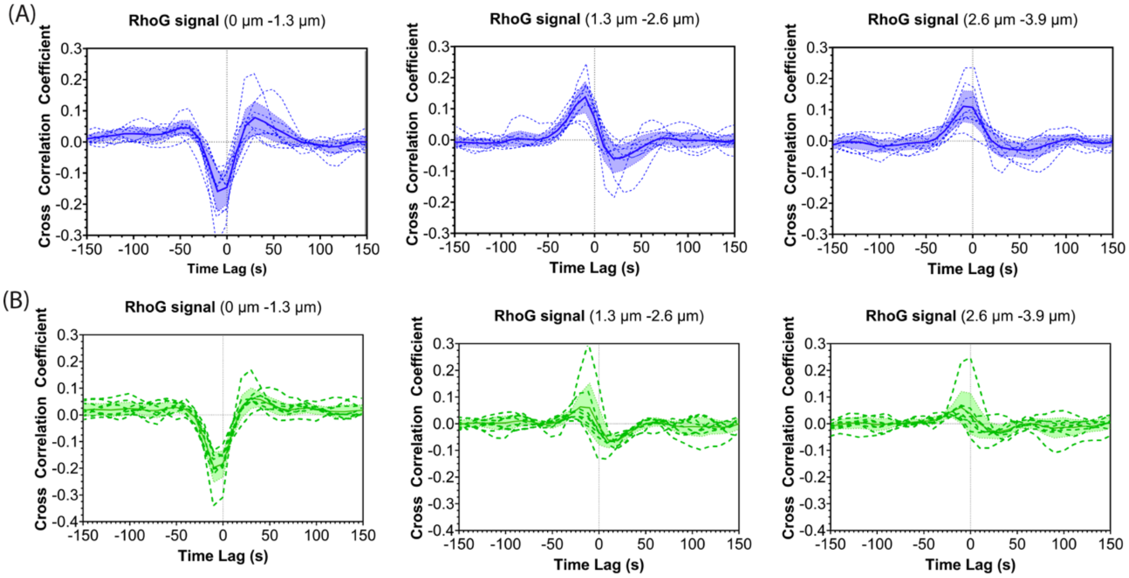


**Fig. S4. Cross-correlation of RhoG activity with cell edge**. Temporal cross-correlations between RhoG biosensor activity and associated edge velocity averaged over the windows in layers 1 – 3 for **(A)** RhoG FLARE.dc (n = 9 cells) and **(B)** Red RhoG FLARE.dc (n = 8 cells). Bold line in each panel shows the averaged cross-correlation over all cells with shaded band indicating the 95% confidence interval. The averaged correlations in layers 1 and 2 from (A) are reproduced in Fig. 1D.

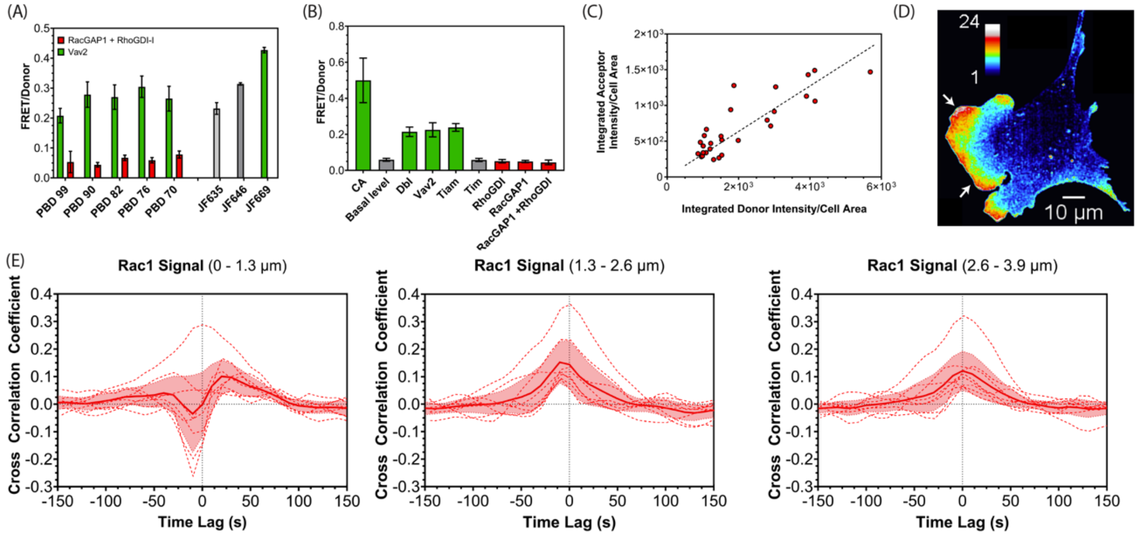


**Fig. S5. Red shifted Rac1 FRET biosensor**. (A) Optimization of FRET/donor ratio by varying the length of the affinity reagent (PBD) and the acceptor dye (error bars are SD, n≥3). Rac1 with constitutively Active (CA, Q61L, Green) or Dominant Negative (DN, T17N, red) mutations were compared to maximize the FRET/Donor ratio. (B) FRET efficiency of the biosensor alone (Basal Level, gray) or when exposed to upstream activators (green), inhibitors (red), upstream regulators not specific for the tested GTPase (gray), and activating point mutations (green, CA) (error bars SD, n = 3). (C) Plot of Donor (mScarlet) and Acceptor (JF669 bound to HaloTag) expression for MEFs stably expressing the dual-chain FRET RhoG biosensor. The ratio of donor to acceptor remains constant across a range of expression levels. Each dot represents an individual cell. (D) Activity maps of the Rac1 biosensor in a randomly moving MEF. Pseudocolor scales indicate ratio values relative to the lowest values. (E) Cross correlation of the Rac1 biosensor activity and edge velocity, showing the lag between changes in protrusion velocity and GTPase activation (n = 8, 95% CI).

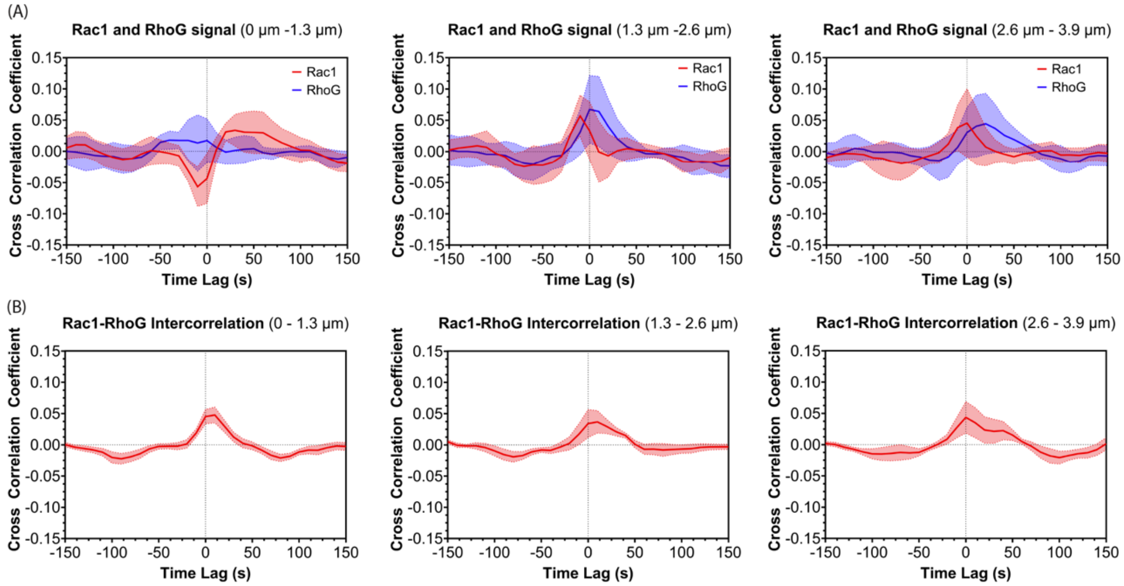


**Fig. S6. Rac1 and RhoG biosensors imaged in the same cell.** (A) Cross-correlation between GTPase activity (RhoG, blue; Rac1, red) and edge velocity for all 3 layers (95% CI, n=9). (B) Intercorrelation of RhoG and Rac1 activities for all 3 layers; to filter the substantial noise only windows were used where the cross-correlation of both biosensor activities and edge velocity was above a threshold of 0.1 (95% CI, n=9).

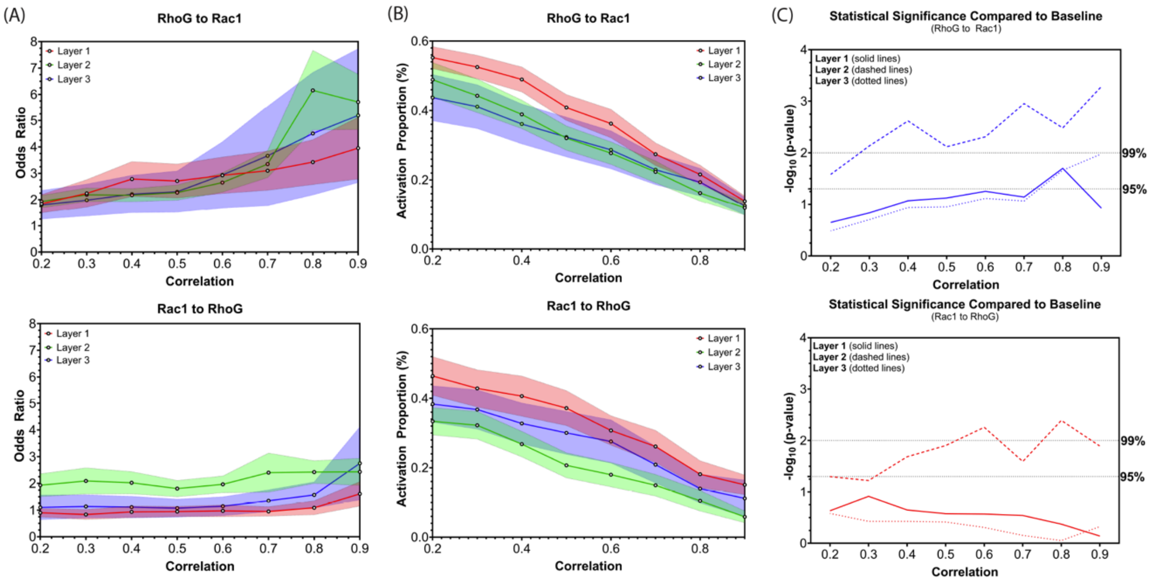


**Fig. S7. Crosstalk Analysis of Rac1 and RhoG activation.** (A) Odds ratio of RhoG activation events leading to Rac1 activation (top) or vice-versa (bottom) as a function of correlation between GTPase activation (error bands 95% CI, n=10). (B) Activation proportion (percentage) of RhoG activation events leading to Rac1 activation (top) or vice-versa (bottom) as a function of correlation between GTPase activation (error bands are 95% CI, n=10). (C) T-test comparison of baseline odds ratio (i.e. 1) with RhoG activation events leading to Rac1 activation (top) or vice-versa (bottom) as a function of correlation of GTPase activation.

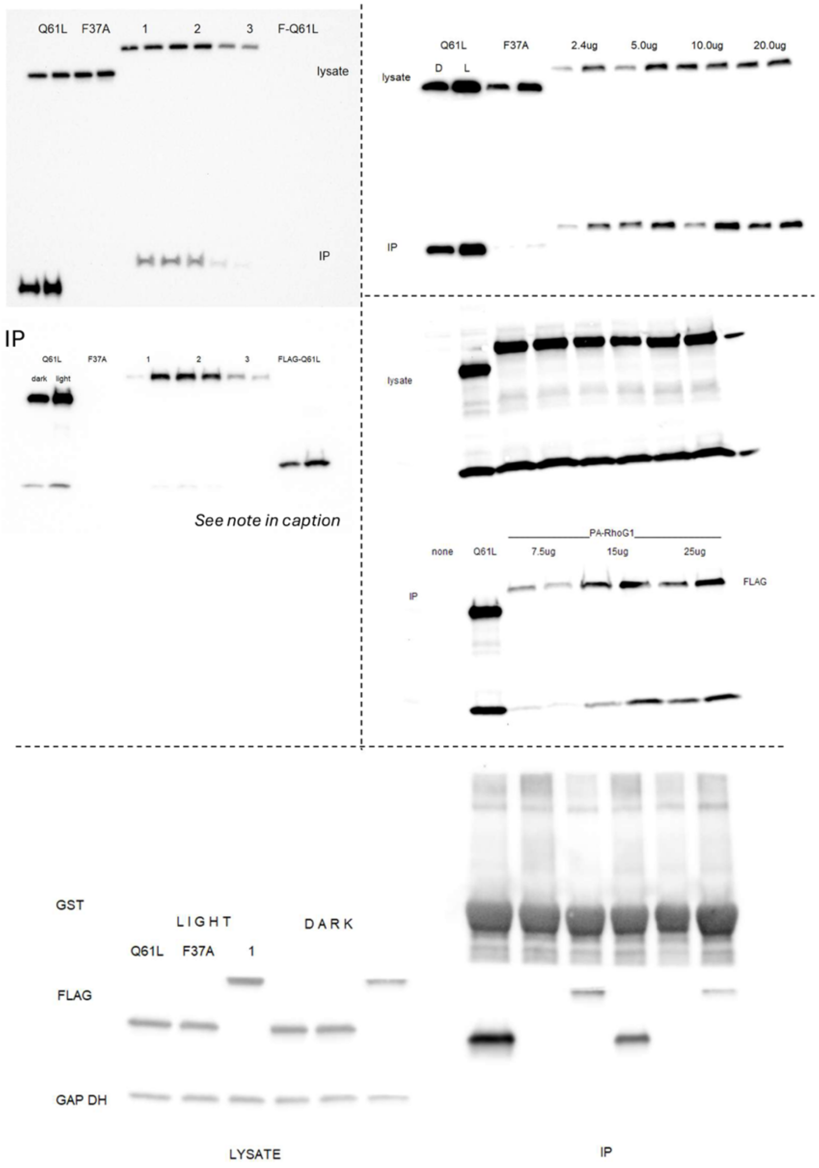


**Fig. S8. Western Blots from PA-RhoG (light and dark conditions) and RhoG from immunoprecipitation (IP) with ELMO beads.** Data used for Figure 2B in the main text. Crops of the first gel (top left) are presented in Fig 1B of the main text. The top image shows both lysate and IP stained. Because the signal was too weak for the IP, the samples were reblotted and re-imaged (bottom).

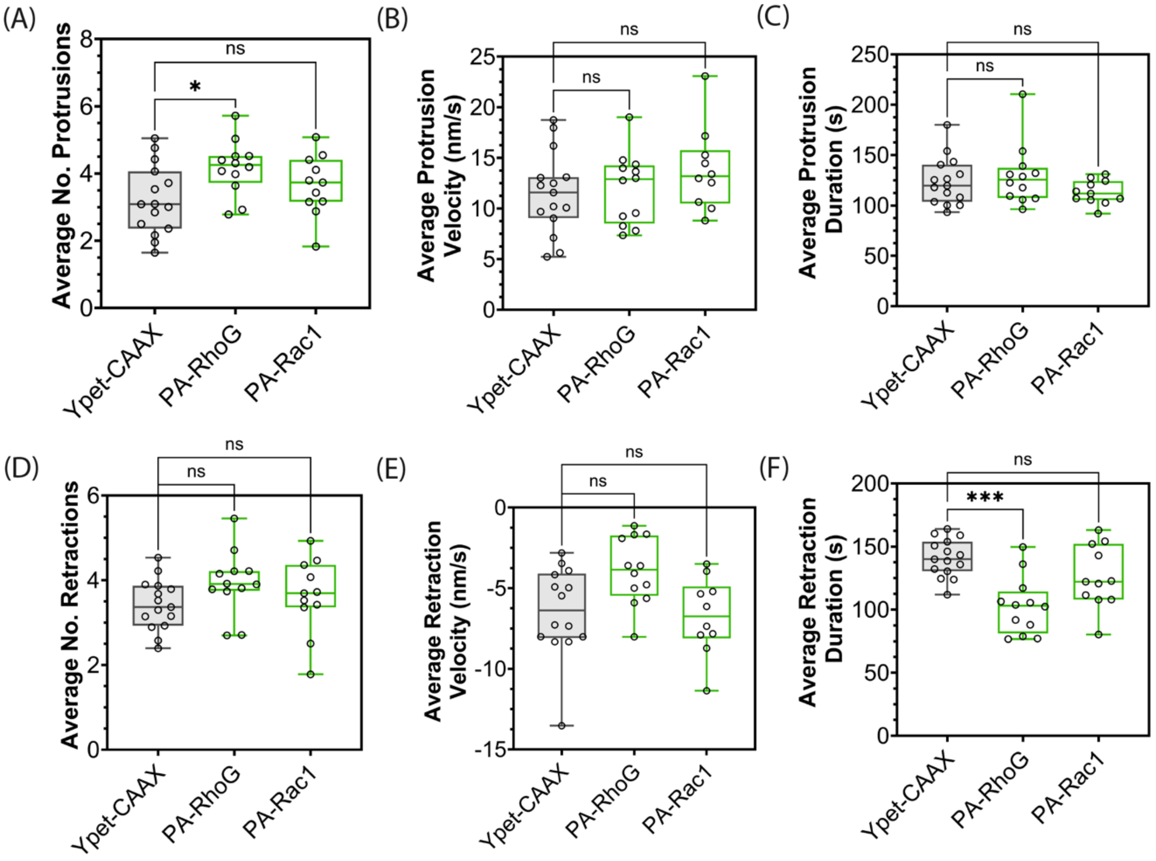


**Fig. S9. Protrusion and retraction parameters of MEFs expressing PA-Rac1 or PA-RhoG in the absence of photoactivation.** MEFs expressing Ypet-CAAX (n=15, control), PA-RhoG (n=12) or PA-Rac1 (n=11). PA-RhoG expression resulted in a statistically significant decrease in the duration of retractions. Dunn's Test, one-way ANOVA non-parametric test (Kruskal–Wallis) used for statistical analysis; * - $p<0.05$, ** - $p<0.01$, *** - $p<0.001$.

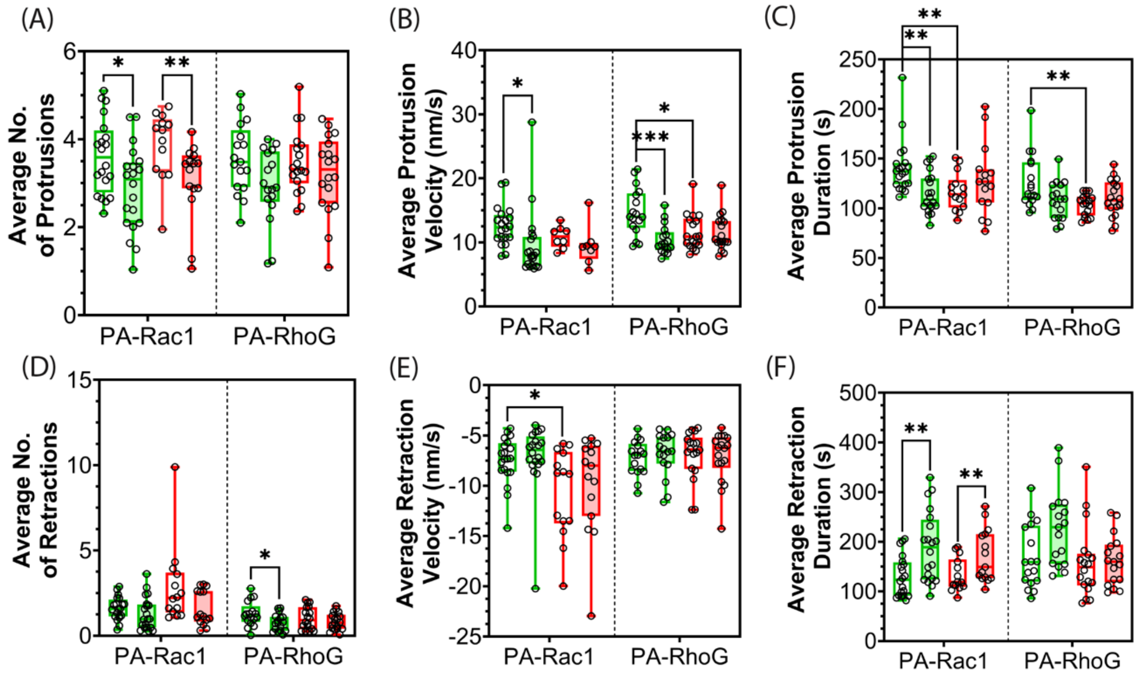


**Fig. S10. Effects of localized PA-Rac1 or PA-RhoG photoactivation on protrusion parameters absent of normalization for inter-cell variability.** Number, velocity and duration of protrusions (A-C) and retractions (D-F) were quantified inside (open boxplots) or outside (shaded boxplots) the photoactivation region for each individual cell. Inside photoactivation region was defined as the parts of the cell within a 346 $\mu m^2$ area (~80-pixel diameter) from the irradiation spot. Optogenetic proteins with wild type LOV are shown in green (n=20 for Rac1, n=17 for RhoG), optogenetic tools with LOV dark mutant are depicted in red (n=13 for Rac1, n=18 for RhoG). See Materials and Methods regading the region selection. Statistical analysis using Dunn's Test, one-way ANOVA non-parametric test (Kruskal–Wallis). * - p<0.05, ** - p<0.01, *** - p<0.005. See Supplementary Text 1 for more information.

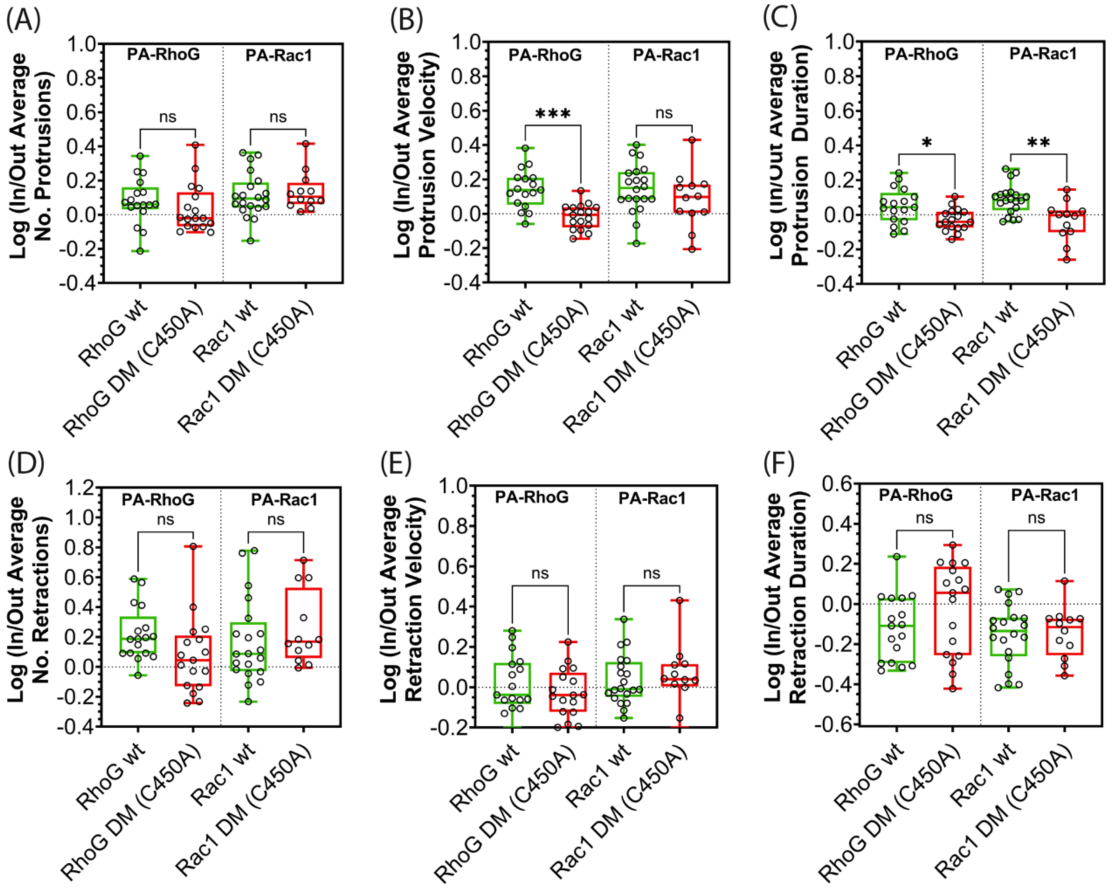


**Fig. S11. Protrusion and retraction dynamics of MEFs upon exposure to localized photoactivation of PA-Rac1 or PA-RhoG normalized for inter-cell variability.** Number, velocity and duration of protrusion (A-C) and retraction (D-F) events for MEFs locally irradiated with blue light expressing optogenetic tools with LOV-wt (green, n=20 for Rac1, n=17 for RhoG), optogenetic tools with LOV-DM (red, n=12 for Rac1, n=17 for RhoG). See Materials and Methods for more details on region selection. Statistical analysis using Dunn's Test, one-way ANOVA non-parametric test (Kruskal–Wallis). * - $p<0.05$, ** - $p<0.01$, *** - $p<0.005$. See Supplementary Text 1 for more details.

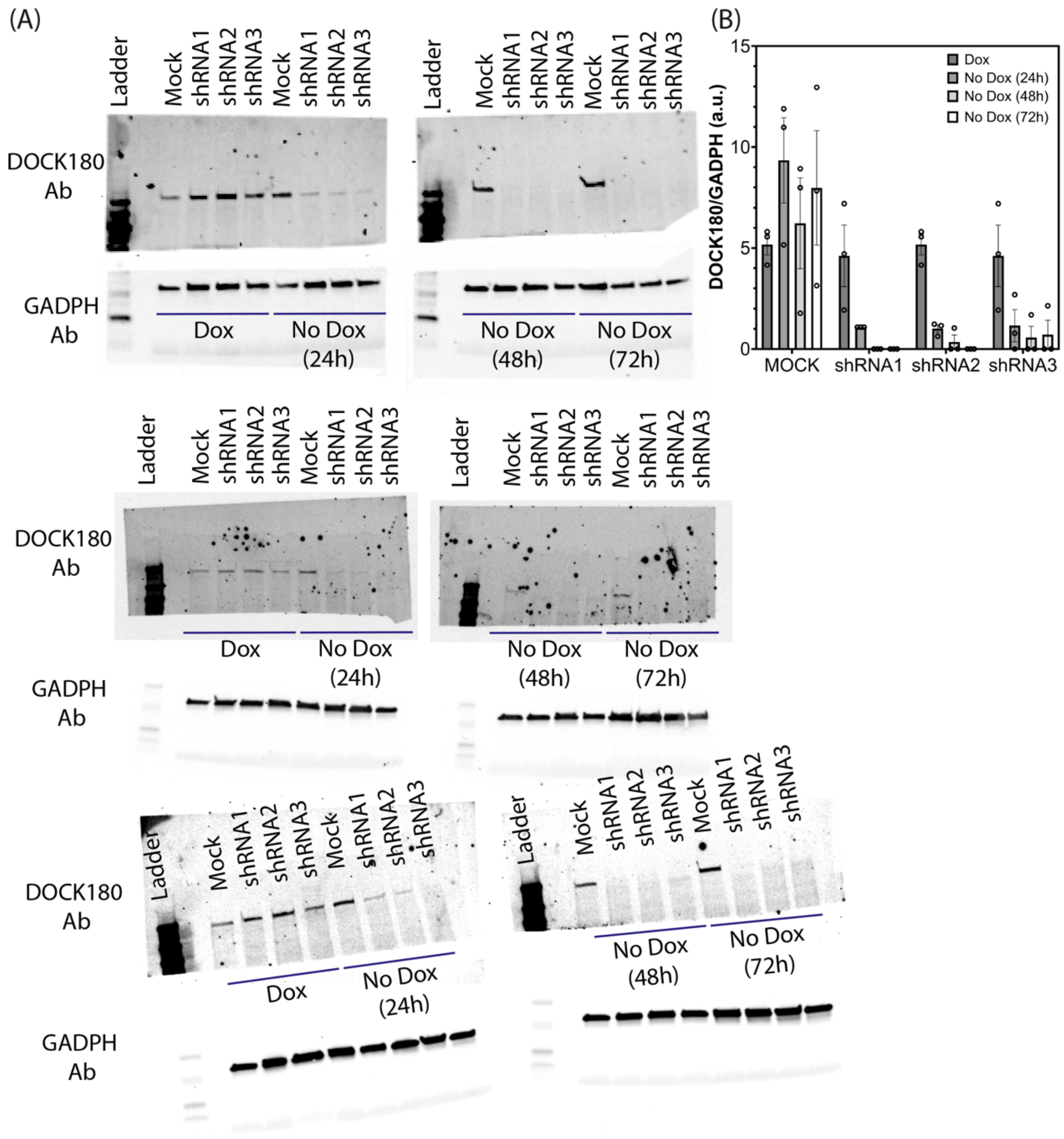


**Fig. S12. shRNAs for DOCK180. (A)** Western blots for DOCK 180 and GADPH (control for number of cells). DOCK180 expression was measured in MEFs stably expressing mock or one of three tested shRNAs in the presence of Doxycycline (Dox, inhibitor of shRNA expression), 24, 48 or 72 hours after Dox removal. **(B)** Quantification of Western blots with DOCK180 expression normalized for GADPH expression.

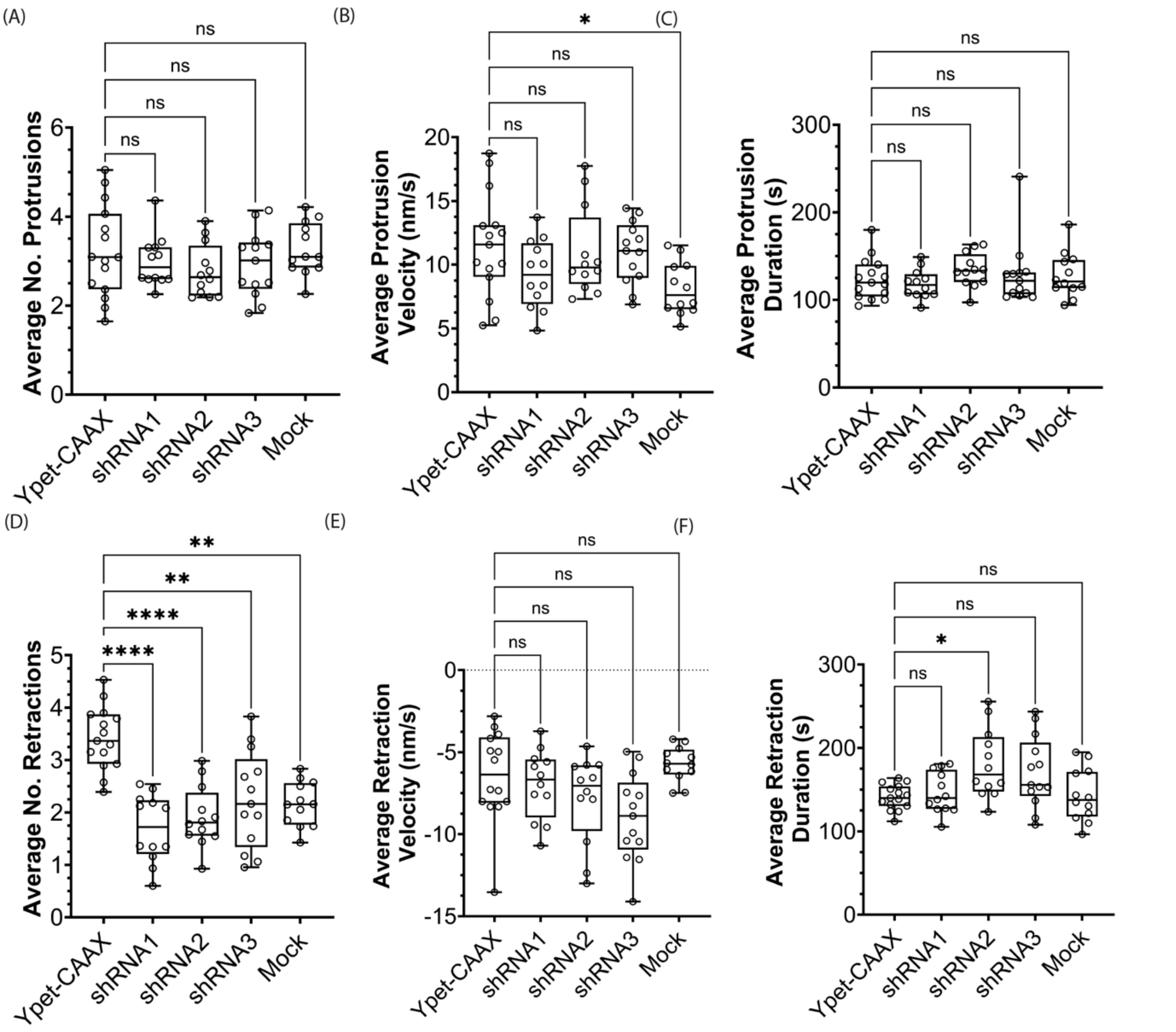


**Fig. S13. Protrusion and retraction parameters of MEFs expressing different shRNAs for DOCK180.** MEFs expressing Ypet-CAAX (n=15, control), DOCK180 shRNA1 (n=12), DOCK180 shRNA2 (n=12), DOCK180 shRNA3 (n=13) or DOCK180 mock shRNA (n=12). Mock showed a statistically significant reduction in the number of retractions compared to control. Mock and the 3 shRNAs for DOCK180 were statistically identical. This indicated that knockdown of DOCK180 did not perturb protrusion and retraction parameters.

### Supplementary Movie Titles and Captions

**Movie S1. RhoG cellular distribution:** Mouse embryonic fibroblast (MEF) randomly moving on fibronectin, stably expressing fluorescently labelled RhoG. RhoG in seen primarily in the perinuclear region, Golgi apparatus and trafficking vesicles.

**Movie S2. RhoG spatio-temporal activity**: Mouse embryonic fibroblast (MEF) randomly moving on fibronectin stably expressing RhoG FLARE.dc.

**Movie S3. Imaging the activities of Rac1 and RhoG simultaneously:** Mouse embryonic fibroblast on fibronectin, stably expressing RhoG FLARE.dc and Far-Red Rac1 FLARE.dc.

**Movie S4. Photoactivation of PA-RhoG is sufficient to induce protrusions:** Mouse embryonic fibroblast (MEF) stably expressing PA-RhoG labelled with iRFP720. Protrusions are induced as a function of localized irradiation in 3 regions.

**Movie S5. Photoactivation of PA-RhoG while imaging Rac1:** Mouse embryonic fibroblast (MEF) stably expressing PA-RhoG labelled with iRFP720 and and Far-Red Rac1 FLARE.dc.
